# The structural and mechanistic bases for the viral resistance to allosteric HIV-1 integrase inhibitor pirmitegravir

**DOI:** 10.1101/2024.01.26.577387

**Authors:** Tung Dinh, Zahira Tber, Juan S. Rey, Seema Mengshetti, Arun S. Annamalai, Reed Haney, Lorenzo Briganti, Franck Amblard, James R. Fuchs, Peter Cherepanov, Kyungjin Kim, Raymond F. Schinazi, Juan R. Perilla, Baek Kim, Mamuka Kvaratskhelia

**Author notes:** Contributed equally. **Conflict of interest statement:** The authors declare a conflict of interest. Kyungjin Kim is the Chief Executive Officer of ST Pharm Co. Ltd. No other authors declare a potential conflict of interest.

## Abstract

Allosteric HIV-1 integrase (IN) inhibitors (ALLINIs) are investigational antiretroviral agents which potently impair virion maturation by inducing hyper-multimerization of IN and inhibiting its interaction with viral genomic RNA. The pyrrolopyridine-based ALLINI pirmitegravir (PIR) has recently advanced into Phase 2a clinical trials. Previous cell culture based viral breakthrough assays identified the HIV-1_(Y99H/A128T IN)_ variant that confers substantial resistance to this inhibitor. Here, we have elucidated the unexpected mechanism of viral resistance to PIR. While both Tyr99 and Ala128 are positioned within the inhibitor binding V-shaped cavity at the IN catalytic core domain (CCD) dimer interface, the Y99H/A128T IN mutations did not substantially affect direct binding of PIR to the CCD dimer or functional oligomerization of full-length IN. Instead, the drug-resistant mutations introduced a steric hindrance at the inhibitor mediated interface between CCD and C-terminal domain (CTD) and compromised CTD binding to the CCD_Y99H/A128T_ + PIR complex. Consequently, full-length IN_Y99H/A128T_ was substantially less susceptible to the PIR induced hyper-multimerization than the WT protein, and HIV-1_(Y99H/A128T IN)_ conferred >150- fold resistance to the inhibitor compared to the WT virus. By rationally modifying PIR we have developed its analog EKC110, which readily induced hyper-multimerization of IN_Y99H/A128T_ *in vitro* and was ∼14-fold more potent against HIV-1_(Y99H/A128T IN)_ than the parent inhibitor. These findings suggest a path for developing improved PIR chemotypes with a higher barrier to resistance for their potential clinical use.

**IMPORTANCE:** Antiretroviral therapies save the lives of millions of people living with HIV (PLWH). However, evolution of multi-drug-resistant viral phenotypes is a major clinical problem, and there are limited or no treatment options for heavily treatment-experienced PLWH. Allosteric HIV-1 integrase inhibitors (ALLINIs) are a novel class of antiretroviral compounds which work by a unique mechanism of binding to the non-catalytic site on the viral protein and inducing aberrant integrase multimerization. Accordingly, ALLINIs potently inhibit both wild type HIV-1 and all drug-resistant viral phenotypes that have so far emerged against currently used therapies. Pirmitegravir, a highly potent and safe investigational ALLINI, is currently advancing through clinical trials. Here we have elucidated structural and mechanistic bases behind the emergence of HIV-1 integrase mutations in infected cell that confer resistance to pirmitegravir. In turn, our findings allowed us to rationally develop an improved ALLINI with substantially enhanced potency against the pirmitegravir resistant virus.

## INTRODUCTION

HIV-1 integrase (IN) is essential for two distinct steps in the virus lifecycle: i) its enzymatic activities are needed for integration of the double-stranded viral complementary DNA into host cell chromosome; ii) during virion morphogenesis IN binds to the viral RNA genome (vRNA) to ensure proper localization of ribonucleoprotein complexes within the mature capsid. The catalytic activity of IN has been exploited as a therapeutic target, and the IN strand transfer inhibitors (INSTIs) have been successfully used to treat people living with HIV.

More recently, allosteric HIV-1 integrase inhibitors (ALLINIs), which target a non-catalytic site on IN, have been developed (1–10). The principal mode of action of ALLINIs is to induce aberrant or hyper-multimerization of the retroviral protein, which is detrimental for both catalytic and non-catalytic functions of IN during early and late steps of HIV-1 replication (3, 11–15). However, in cell culture ALLINIs much more potently inhibit proper virion maturation than HIV-1 integration (3, 7, 8, 16–18). The cellular cofactor LEDGF/p75, which mediates integration of HIV-1 into active transcription units (19–21), binds IN at the same non-catalytic dimer interface which is targeted by ALLINIs (22, 23). Accordingly, the competitive interplay between nuclear LEDGF/p75 and ALLINIs during HIV-1 integration substantially reduces the potency of these inhibitors in target cells (24, 25). Overexpression of LEDGF/p75 further decreases ALLINI EC_50_ values whereas the LEDGF/p75 depletion substantially enhances the potency of these inhibitors during early steps of infection (24). By contrast, during virion morphogenesis ALLINIs readily induce hyper-multimerization of IN and impair its binding to viral RNA (12). Consequently, the virions produced in the presence of ALLINIs have ribonucleoprotein complexes mislocalized outside of the protective capsid and are non-infectious (3, 7, 12, 16–18, 26–29).

ALLINIs typically contain core aromatic scaffolds, which are flanked by the key pharmacophore carboxylic acid, the *tert*-butoxyl moiety and halogenated bulky aromatic rings. These inhibitors are anchored to the V-shaped cavity at the IN catalytic core domain (CCD) dimer through an extensive network of hydrogen bonds and hydrophobic interactions (1–3, 7). Biochemical assays have revealed that in addition to CCD, the C-terminal domain (CTD) of IN is crucial for ALLINI induced aberrant protein multimerization (30). More recent X-ray crystallographic studies have elucidated the structural basis for ALLINI induced aberrant IN multimerization (6, 31–33). These inhibitors induce head to tail interactions between CCD-CCD of one dimer and the CTD of another dimer, which lead to the uncontrolled hyper-multimerization of IN, thereby resulting in non-functional protein polymers. The invariant CTD residues engage with both CCD and the inhibitor to stabilize the CCD-ALLINI-CTD interface. Because ALLINIs are sandwiched between CCD and CTD, these inhibitors exhibit a substantially lower *K_off_* rate and a higher affinity (*K_D_*) for the CCD-ALLINI-CTD vs CCD-ALLINI complex (32).

Over the past decade multiple ALLINI chemotypes with different core aromatic ring structures have been developed (1–8). Of these, pirmitegravir (Fig 1A), which contains the core pyrrolopyridine ring, has recently advanced into Phase 2a clinical trials. The cell culture-based viral breakthrough assays have identified IN mutations that arose under the selective pressure of PIR (5). Initial evolution of the HIV-1_(Y99H IN)_ phenotype was followed by the emergence of the HIV-1_(Y99H/A128T IN)_ variant at higher PIR concentrations. Here, we have investigated structural and mechanistic bases for the viral resistance to PIR. Surprisingly, we found that even though Tyr99 and Ala128 are positioned within the V-shaped cavity at the CCD dimer, the Y99H/A128T IN changes did not substantially affect direct binding of PIR to CCD. Instead, the steric hindrance induced by the resistant mutations prevented the CTD binding to the CCD_Y99H/A128T_ + PIR. By exploiting these structural findings, we have rationally developed an improved PIR analog EKC110, which exhibited ∼14-fold higher potency against HIV-1_(Y99H/A128T IN)_ compared to its parental inhibitor.

**FIG 1.**
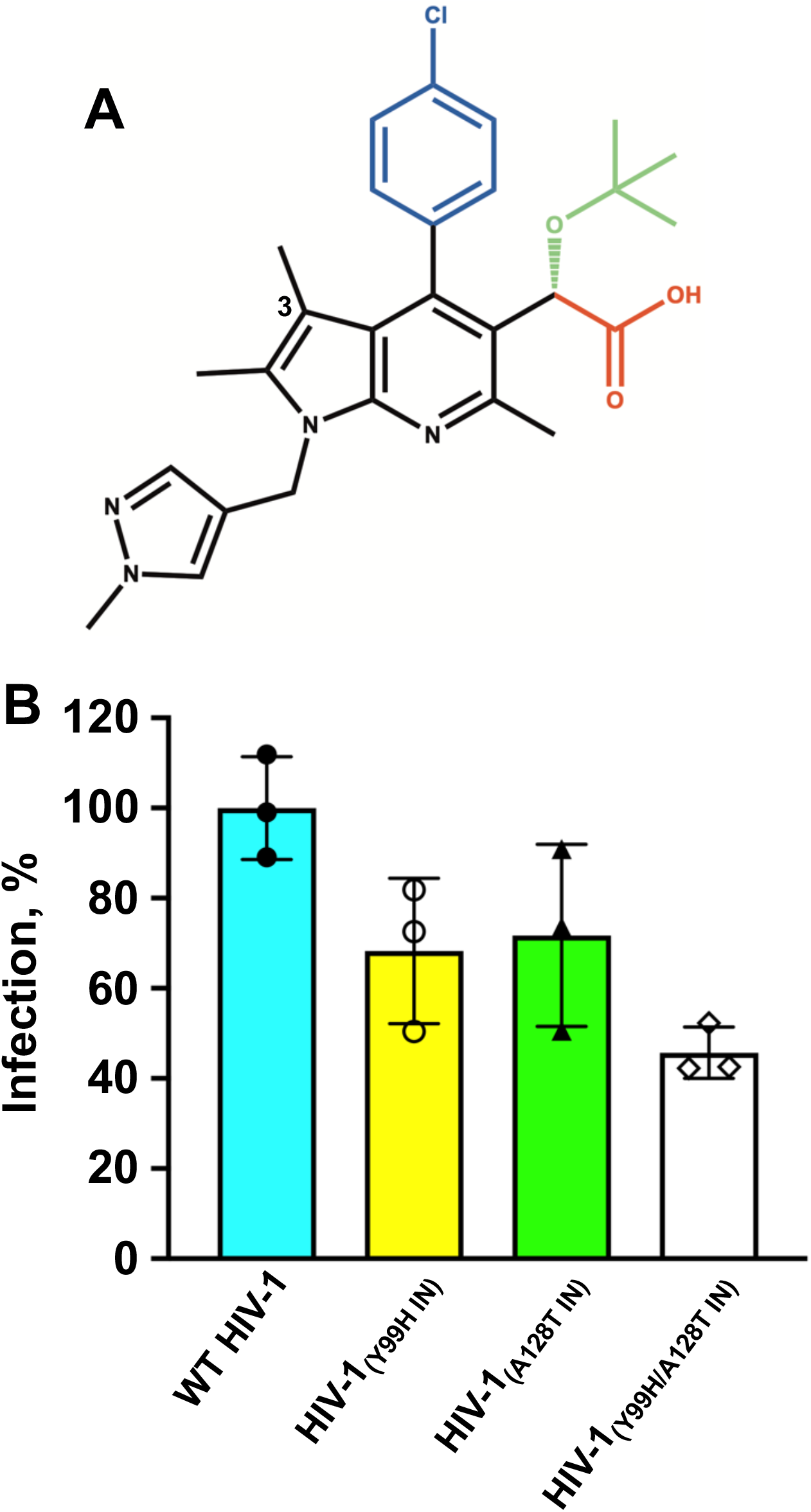
(A) The chemical structure of PIR. The separate functional groups are color-coded: carboxylate (red); *tert*-butoxyl (green); chlorophenyl (blue); core pyrrolopyridine and methylpyrazole rings (black). The 3-methyl group on the pyrrolopyridine ring is indicated. (B) Infectivity of WT and indicated mutant viruses.

## RESULTS

### Effects of Y99H, A128T and Y99H/A128T IN mutations on antiviral activity of PIR

We evaluated how Y99H, A128T and Y99H/A128T IN substitutions affect HIV-1 replication. The virus production from HEK293T cells transfected with full-length NL4.3 plasmids was not detectably influenced by these amino acid changes (Fig. S1). Infectivity of single mutant viruses HIV-1_(Y99H IN)_ and HIV-1_(A128T IN)_ in TZM-bl cells were reduced by ∼30%, whereas HIV-1_(Y99H/A128T IN)_ was ∼55% less infectious compared to WT HIV-1 (Fig. 1B). The antiviral assays performed with PIR revealed that HIV-1_(Y99H IN)_ conferred relatively modest (∼4-fold) resistance to the inhibitor, whereas larger reductions in the inhibitor potency were observed with respect to HIV-1_A128T IN_ (∼13-fold) and HIV-1_Y99H/A128T IN_ (>150-fold) compared to their WT counterpart (Table 1 and Fig. S2).

**Table 1.** Antiviral activities of PIR against WT and indicated mutant viruses.

| HIV-1<sub>NL4.3</sub> | PIR, nM |
| --- | --- |
| WT | 10.4 ± 0.4 |
| Y99H | 39.9 ± 4.8 |
| A128T | 132.9 ± 31.9 |
| Y99H/A128T | 1559.8 ± 62.2 |

### Biochemical mechanisms of the IN_Y99H/A128T_ resistance to PIR

Our biochemical assays have focused on elucidating the underlying mechanism for the major drug resistant IN_Y99H/A128T_ protein. Introduction of the Y99H/A128T changes in the full-length IN did not substantially affect its oligomeric state (Fig. S3). However, marked differences between WT and Y99H/A128T INs were evident upon addition of PIR (Fig. 2). Dynamic light scattering (DLS) assays demonstrated that PIR rapidly (within 1 min) induced hyper-multimerization of WT IN yielding aberrant protein aggregates with hydrodynamic radii of >400 nm (Fig. 2A). In contrast, no protein aggregates were observed after 10 min of PIR addition to IN_Y99H/A128T_ (Fig 2B). Instead, initial protein aggregation with particle sizes of <100 nm was detected only after 15 min of incubation of PIR with the mutant protein (Fig. 2B).

**FIG 2.**
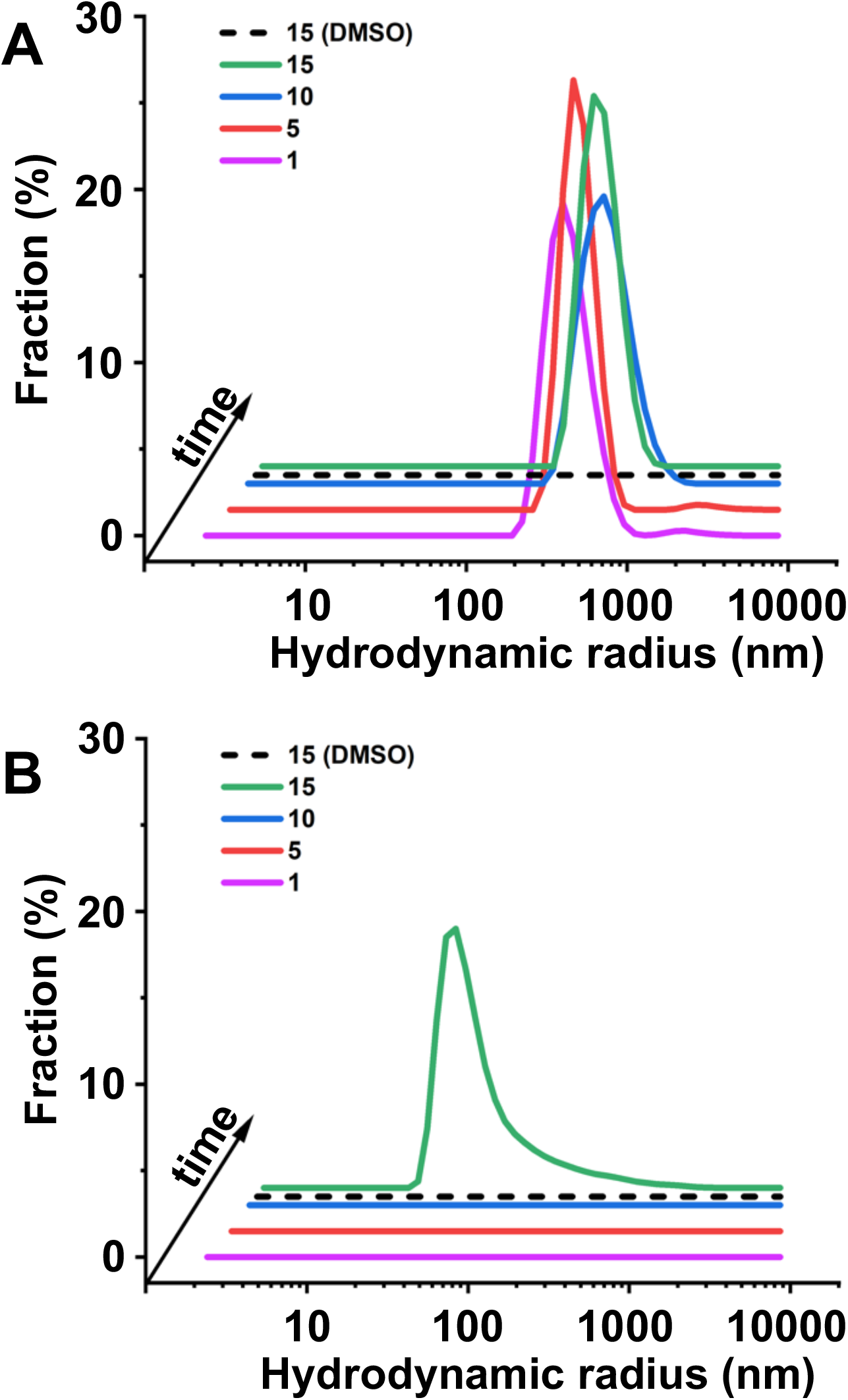
DLS analysis of PIR induced aberrant IN multimerization. 500 nM PIR was added to 200 nM full-length WT IN (A) or IN_Y99H/A128T_ (B) and DLS signals were recorded at indicated times (1-15 min). DMSO controls are shown after incubation of full-length IN proteins for 15 min to indicate that these proteins remained fully soluble in the absence of PIR.

To understand how the Y99H/A128T IN mutations affect the inhibitor binding to the CCD dimer we conducted surface plasmon resonance (SPR) assays. Surprisingly, we observed only slight differences between PIR binding to CCD_Y99H/A128T_ (*K_D_* of ∼77 nM) vs WT CCD (*K_D_* of ∼24 nM) (Fig. 3). Clearly, this modest (∼3-fold) change in the binding affinity does not explain the marked resistance (>150-fold) to PIR conferred by HIV-1_(Y99H/A128T IN)_ compared to the WT virus (Table 1).

**FIG 3.**
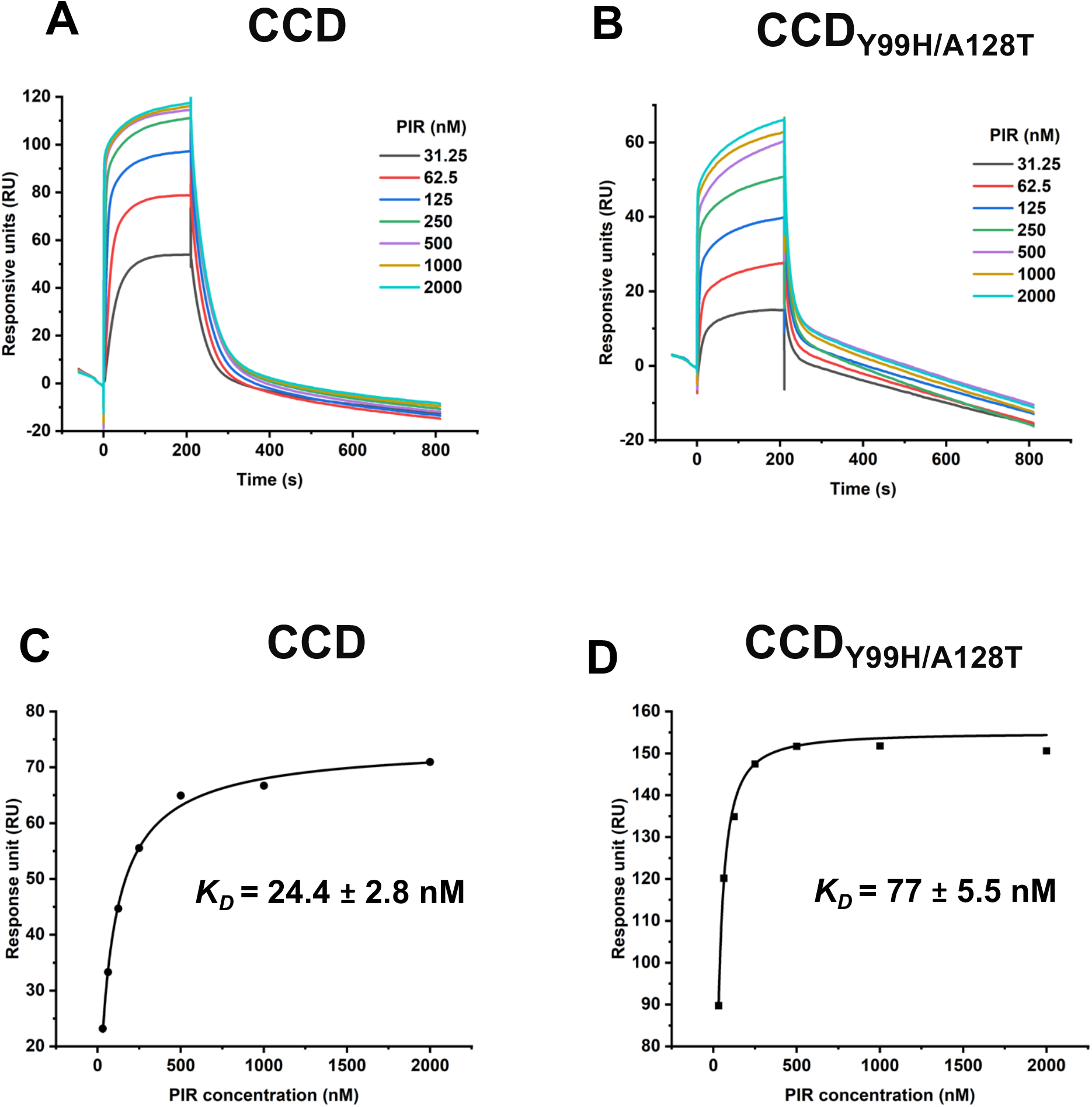
SPR analysis of PIR binding to WT CCD and CCD_Y99H/A128T_. Representative sensorgrams for PIR binding to of WT CCD (A) vs CCD_Y99H/A128T_ (B). PIR concentrations are indicated. The *K_D_* values for PIR + CCD (C) and PIR + CCD_Y99H/A128T_ (D) were determined using the Hill equation.

Upon binding to the V-shaped cavity at the CCD dimer interface, ALLINIs act as molecular glues to recruit CTD (31–33). Therefore, we tested how Y99H/A128T IN mutations affected formation of the CCD-PIR-CTD complex. For this, we have developed an affinity pull-down assay to capture the CTD specifically bound to the His_6_-CCD in the complex with PIR. The results in Fig. 4 demonstrate that CTD was selectively pulled-down by His_6_-CCD only in the presence, but not in the absence, of PIR (Fig. 4, compare lane 9 with 6). In sharp contrast from WT CCD, His_6_-CCD_Y99H/A128T_ failed to bind to CTD in the absence or presence of PIR (Fig. 4, lanes 7 and 10).

**FIG 4.**
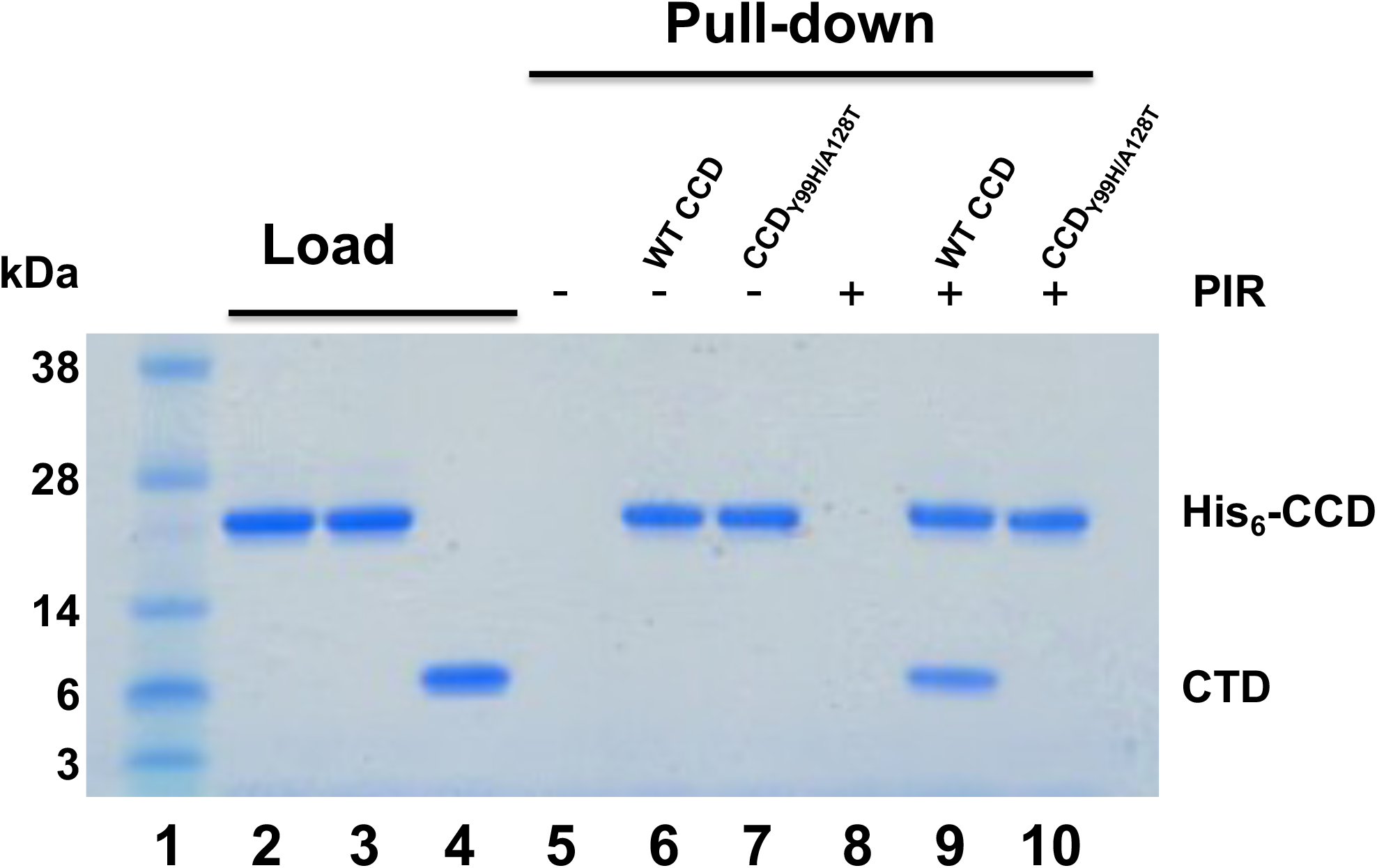
Affinity pull-down assays to probe PIR induced CCD-CTD interactions. Lane 1: molecular weight markers; Lanes 2 – 4: loads of His_6_-CCD (lane 2), His_6_-CCD_Y99H/A128T_ (lane 3), and tag-less CTD (lane 4); Lanes 5 - 7: affinity pull-down using Ni beads of CTD alone (lane 5, control), His_6_-CCD + CTD (lane 6), His_6_-His_6_-CCD_Y99H/A128T_ + CTD (lane 7) in the absence of PIR; Lanes 8 - 10: affinity pull-down using Ni beads of CTD + PIR (lane 8, control), His_6_-CCD + CTD (lane 9), His_6_-His_6_-CCD_Y99H/A128T_ + PIR + CTD (lane 10).

Taken together, our biochemical studies indicate that the Y99H/A128T IN changes do not substantially affect direct binding of PIR to its cognate V-shaped cavity at the CCD dimer. Instead, the Y99H/A128T mutations strongly interfere with CTD binding to the CCD + PIR complex. Consequently, IN_Y99H/A128T_ confers the marked resistance with respect to the ability of PIR to induce aberrant IN multimerization.

### The structural basis for the IN_Y99H/A128T_ resistance to PIR

We have solved X-ray structures of PIR bound to both WT CCD and CCD_Y99H/A128T_ (Tables S1, S2), which revealed very similar binding of the inhibitor to these proteins (Fig. 5A). Both the aromatic ring of Tyr99 and the imidazole ring of His99 adopt very similar positions as they extend inside the CCD-CCD dimer interface and away from the bound PIR (Fig. 5A). The Ala128 side chain is surface exposed and extends toward the 3-methyl group of the core pyrrolopyridine ring system. Yet, the substitution of Ala128 with the bulkier and polar Thr128 did not seemingly alter the inhibitor positioning in the V-shaped pocket and the distances from the 3-methyl group to the closest C_β_ of Ala128 and Thr128 were nearly identical (∼3.0 Å vs 3.2 Å). Furthermore, the polar group of Thr128 points away from the inhibitor. Taken together, these structural findings agree well with our biochemical results indicating that the Y99H/A128T changes do not substantially affect functional oligomerization of full length IN (Fig. S3) or the direct binding of the inhibitor to the CCD dimer (Fig. 3).

**FIG 5.**
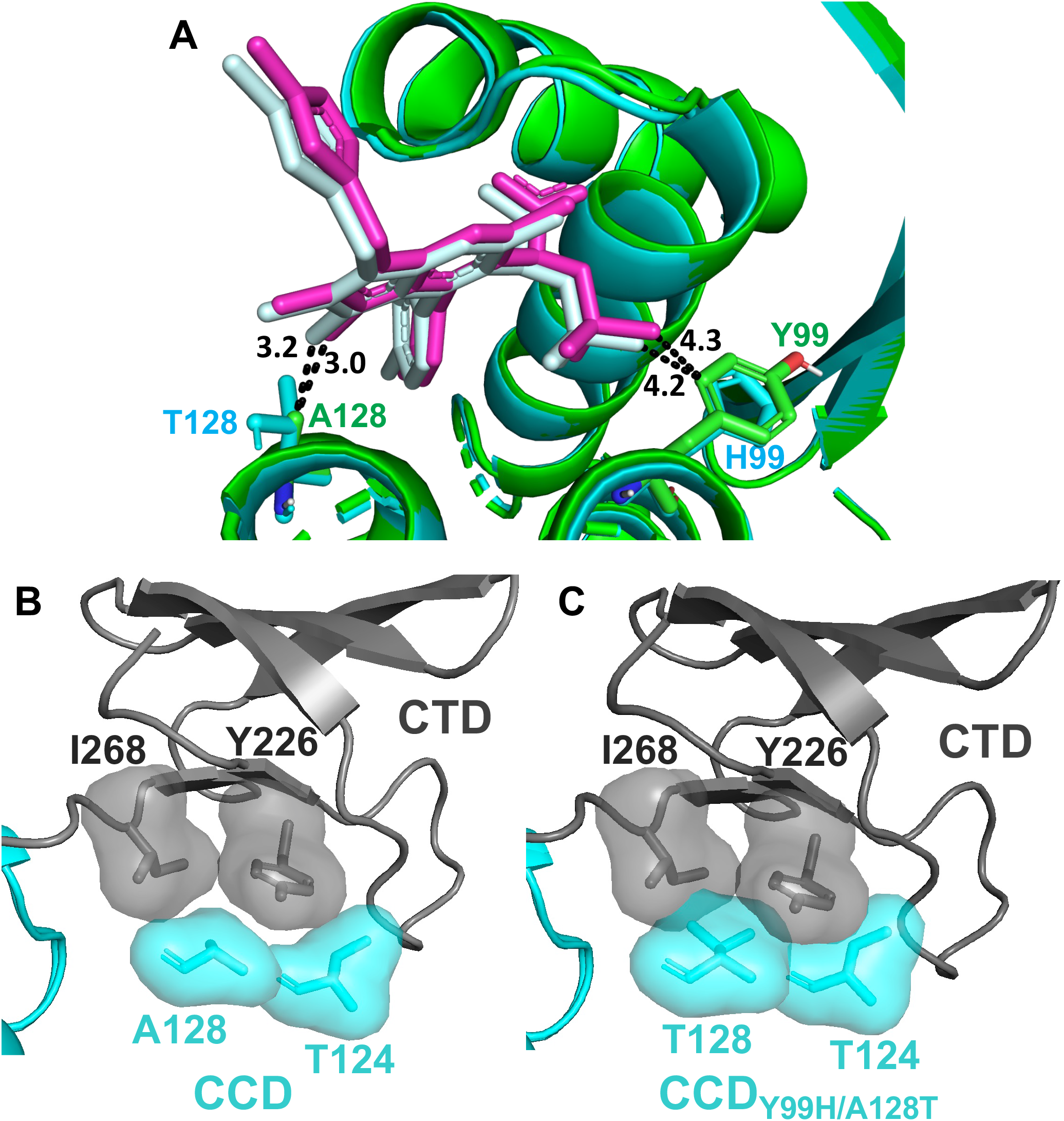
Structural analysis of PIR interactions with CCD vs CCD_Y99H/A128T_. (A) Superimposed crystals structures of WT CCD (green) + PIR (magenta) and CCD_Y99H/A128T_ (cyan) + PIR (pale cyan). Distances were measured between the 3-methyl group of PIR ‘s pyrrolopyridine ring to the closest Cβ on either Ala128 or Thr128, as well as between the closest methyl group on PIR ‘s *tert*-butoxy to either Tyr99 or His99. (B) Van der Waals surface for indicated residues are shown in the structure of WT CTD-CCD + PIR. (C) The structure of CCD_Y99H/A128T_ + PIR superimposed onto the structure of WT CTD-CCD + PIR. Van der Waals surface for indicated residues reveals steric clashes observed by overlapping, shaded surfaces. PIR is not shown for clarity.

Recently, two-domain HIV-1 IN CTD-CCD constructs were developed to study ALLINI-induced CTD-CCD interaction (32). Our efforts to obtain a crystal structure for the CTD-CCD_Y99H/A128T_ + PIR complex have not been successful likely due to the inability of CTD to bind to the CCD_Y99H/A128T_ + PIR complex (Fig. 4). Therefore, to understand how Y99H/A128T mutations affect the CTD binding, we superimposed our crystal structure of CCD_Y99H/A128_ + PIR onto the recently reported structure of the CTD-CCD + PIR (32) (Fig. 5B,C, Fig. S4). Fig. 5B and Fig. S4A show optimal positioning of CCD residues Thr124 and Ala128 with respect to CTD residues Tyr226 and Ile268 at the WT CCD-PIR-CTD interface (32).

Of note, the change of Ala128 to the bulkier and polar Thr128 creates steric hindrance with respect to CTD residues Tyr226 and Ile268 (Fig. 5C, Fig. S4B). Additionally, Thr128 indirectly triggers yet another steric clash between CCD Thr124 and CTD Tyr226. The root cause for this is a hydrogen bond formed between the Thr128 side chain hydroxyl and Thr124 observed in our crystal structure of the CCD_Y99H/A128T_ + PIR complex (Fig. S4B), which in turn repositions Thr128 too close to CTD Tyr226 (Fig. 5C, Fig. S4B). These structural observations are consistent with the experimental results demonstrating that the CCD_Y99H/A128T_ + PIR complex does not effectively interact with the CTD (Fig. 4).

### The development of an improved PIR analog EKC110

From examining crystal structures of PIR bound to WT CCD and CCD_Y99H/A128T_ (Fig. 5A), we noticed that the 3-methyl group of the core pyrrolopyridine ring system extends toward both Ala128 and Thr128, and partly limits PIR accessibility within the V-shaped pocket. We hypothesized that removing the 3-methyl group could enable a modified PIR analog to position itself deeper within the CCD dimer and potentially reduce steric hindrance with respect to the CTD binding to the CCD_Y99H/A128T_ + PIR complex. To test this notion, we have synthesized the PIR analog EKC110 lacking the 3-methyl group (Fig 6A).

**FIG 6.**
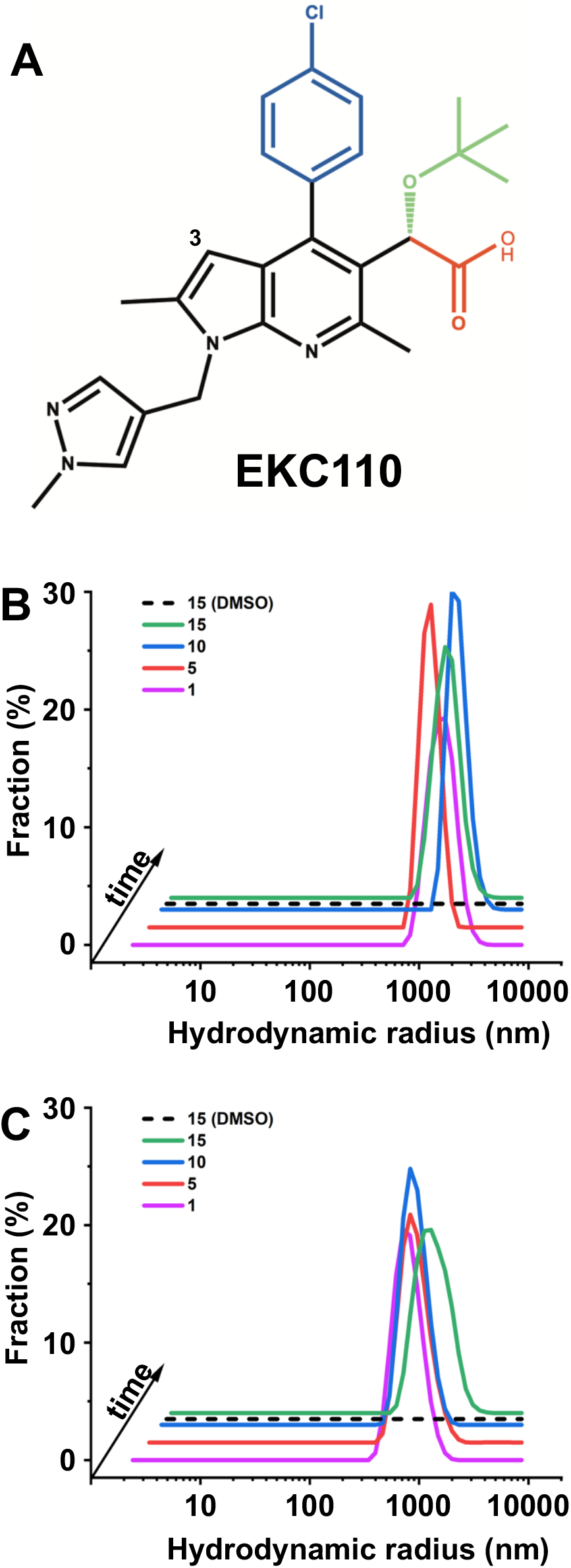
Interactions of EKC110 with HIV-1 IN. (A) The chemical structure of EKC110. The separate functional groups are color-coded: carboxylate (red); *tert*-butoxyl (green); chlorophenyl (blue); core pyrrolopyridine and methylpyrazole rings (black). (B, C) DLS analysis of EKC110 induced aberrant IN multimerization. 500 nM EKC110 was added to 200 nM full-length WT IN (B) or IN_Y99H/A128T_ (C) and DLS signals were recorded at indicated times (1-15 min). DMSO controls are shown after incubation of full-length IN proteins for 15 min to indicate that these proteins remained fully soluble in the absence of EKC110.

Excitingly, EKC110 exhibited ∼14-fold improved potency against HIV-1_(Y99H/A128T IN)_ compared to PIR (Table 2). Furthermore, EKC110 was ∼2-fold more potent than PIR against WT HIV-1. Biochemical assays revealed that unlike PIR (Fig. 2), EKC110 effectively induced aberrant multimerization of both full-length WT and Y99H/A128T INs (Fig. 6B, C).

**Table 2.** Antiviral activities of EKC110 vs PIR.

| HIV-1<sub>NL4.3</sub> | PIR<br>EC<sub>50</sub> (nM) | EKC110<br>EC<sub>50</sub> (nM) | Potency<br>increase, fold |
| --- | --- | --- | --- |
| WT | 10.4 ± 0.4 | 4.5 ± 0.2 | ~2.3 |
| Y99H/A128T | 1559.8 ± 62.2 | 110.7 ± 5.2 | ~14 |

We have solved the X-ray crystal structures of EKC110 bound to WT CCD, CCD_Y99H/A128T_ and WT CTD-CCD (Tables S1, S2, Fig. 7, Fig. S5), whereas the CTD-CCD_Y99H/A128T_ + EKC110 complex did not yield crystals. A comparative analysis of EKC110 with PIR reveals both similarities and notable differences between these inhibitors (Fig. 7, Fig. S5). In common with other members of the ALLINI class of inhibitors, the EKC110 key pharmacophore carboxylic acid establishes bidentate hydrogen bonding with backbone amides of Glu170 and His171 (Fig. S5A). Additionally, the side chain of Thr174 hydrogen bonds with both EKC110 carboxylate and *tert*-butoxy moiety, which is crucial for the high potency of ALLINIs (Fig. S5A). EKC110 positions very similarly within CCD and CCD_Y99H/A128T_ indicating that the drug resistant mutations that confer the marked resistance to PIR do not influence direct binding of EKC110 to the CCD dimer (Fig. S5B).

**FIG 7.**
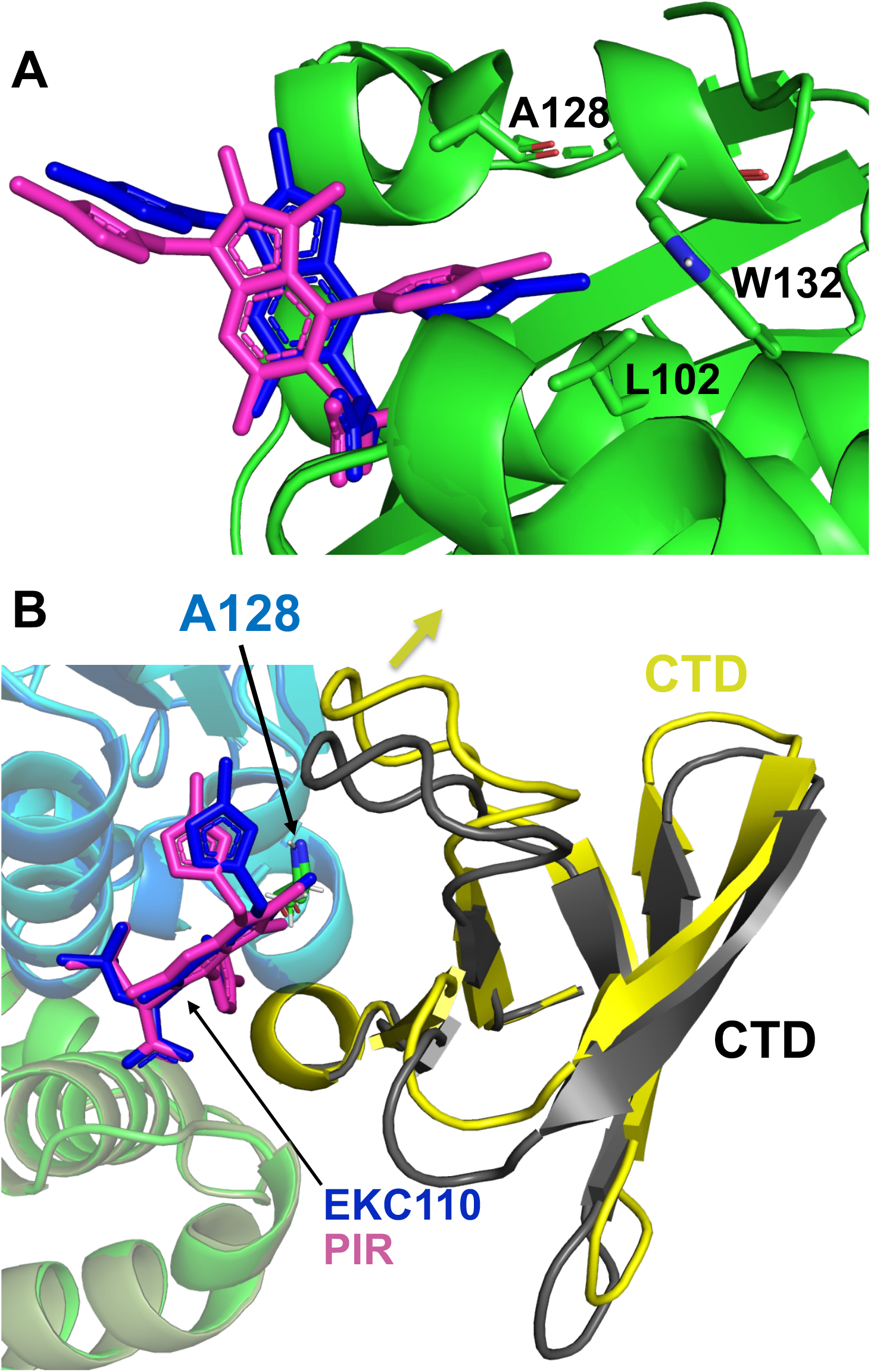
The structural analysis of EKC110 interactions with CCD and CTD-CCD. (A) For comparison the crystal structure of CCD + EKC110 is superimposed onto CCD + PIR, which reveals a noticeable tilt of the EKC110 pyrrolopyridine core toward A128 compared to PIR. C) For comparison the crystal structure of CTD-CCD + EKC110 is superimposed onto CTD-CCD + PIR to show repositioning (yellow arrow) of CTD in the presence of EKC110 compared to PIR. CTDs are shown in yellow and gray in EKC110 + CTD-CCD and PIR + CTD-CCD structures. PIR and EKC110 are in magenta and blue. The side chain of Ala128 in each structure is shown by sticks.

We have observed the following significant differences between EKC110 and PIR binding to either CCD or CTD-CCD (Fig. 7). EKC110 core pyrrolopyridine and methylpyrazole rings are slightly shifted compared to PIR. More specifically, because of the lack of the 3-methyl group, the EKC110 core pyrrolopyridine ring moves closer to and forms hydrophobic interactions with Ala128 (Fig. 7A). Consequently, EKC110 chlorobenzene group extends deeper inside the CCD-CCD dimer cavity toward Trp132 and Leu102 compared to its parental PIR.

Another significant change is a considerable repositioning of CTD at the interfaces mediated by EKC110 vs PIR (Fig. 7B). Specifically, the Cα atom of the CTD residue Tyr226 is shifted away from the Cβ atom of the CCD residues Ala128 by ∼2.5 Å in the complex with EKC110 compared to PIR (Fig. 7B). Collectively, a deeper positioning of EKC110 inside the V-shaped cavity at the CCD dimer coupled with an extended space afforded at the CCD-EKC110-CTD interface compared with the CTD-PIR-CCD complex, could provide structural clues as to why the bulkier Thr128 does not impact EKC110 activity as much as PIR. To further test this notion and better understand differential effects of Y99H/A128T IN mutations on PIR vs EKC110, we performed molecular dynamic (MD) simulations and free energy perturbation (FEP) calculations (see below).

### MD simulations and energetics of PIR and EKC110 interactions with IN_Y99H/A128T_

To quantify the effect of the Y99H/A128T IN mutations on the binding of PIR and EKC110 at the CCD-CTD interface, we performed 1 μs MD simulations (Fig. 8, Fig. S6). Both ALLINIs greatly stabilized interactions between WT CCD and CTD. The average root mean squared deviation (RMSD) were 2.96 ± 0.29 Å and 3.03 ± 0.32 Å for the CCD-PIR-CTD complex; 2.92 ± 0.23Å and 3.00 ± 0.29Å for the CCD-EKC110-CTD complex; and 3.23 ± 0.38Å and 6.62 ± 1.15Å for apo CCD and CTD in the absence of ALLINIs. Of note, during 1 μs MD simulations the CTD domain separated from apo CCD, whereas the CCD-ALLINI-CTD interface remained stable (Fig. S6A) and exhibited an overall decrease in the root mean squared fluctuations (RMSF) across all IN residues (Fig. S6B).

**FIG 8.**
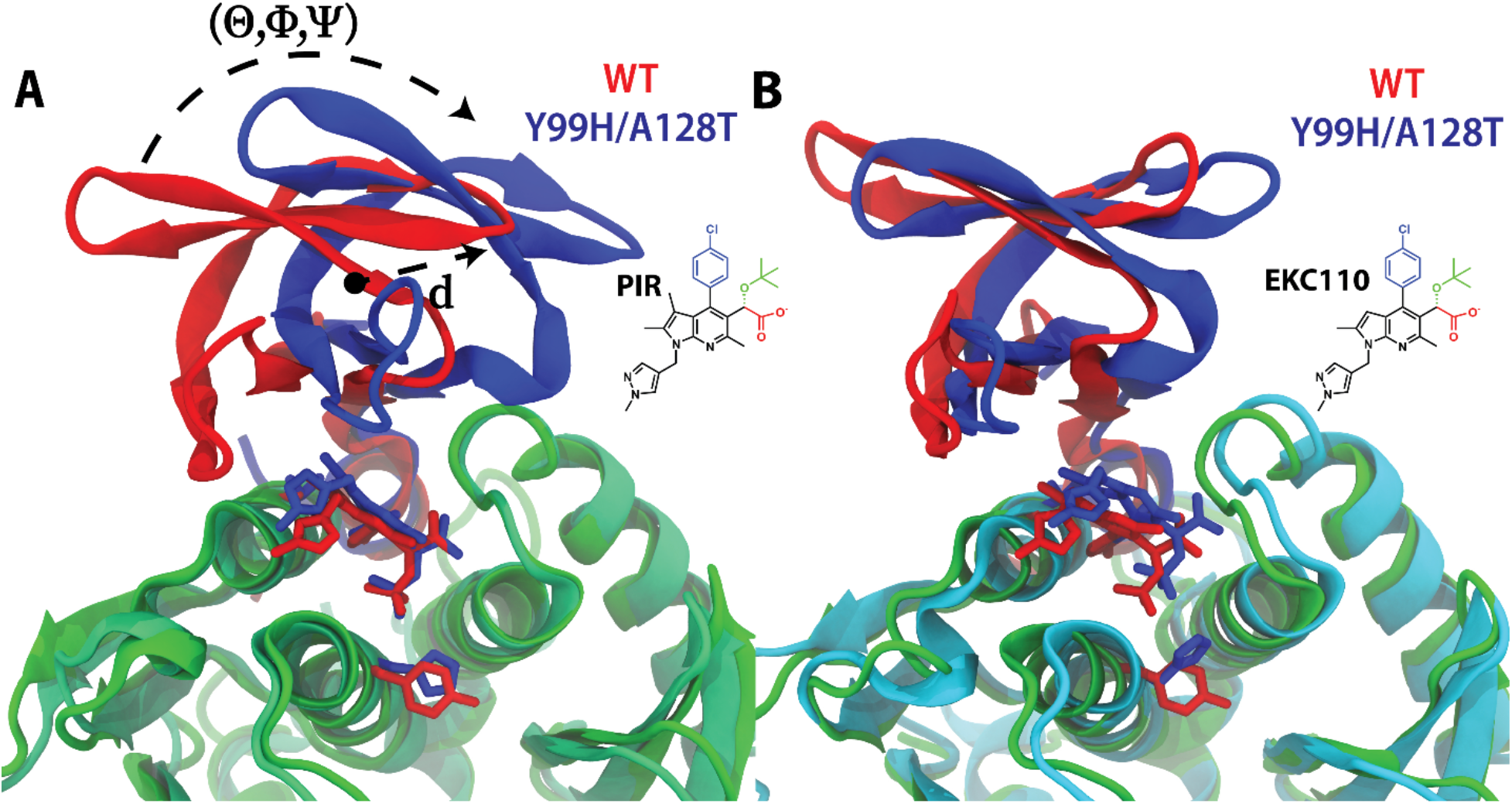
MD simulation of CTD interactions with WT CCD vs CCD_Y99H/A128T_ in the complex with PIR (A) and EKC110 (B). Displacement of the CTD domain from the PIR + CCD_Y99H/A128T_ complex is measured with a center-of-mass displacement of d=4.85Å and rotations along the principal axes of inertia of θ=26.66°, Φ=23.84° and Ψ=11.81°. WT CCD and CCD_Y99H/A128T_ are colored cyan and green, respectively. CTDs interacting with WT CCD and CCD_Y99H/A128T_ are colored red and blue, respectively.

While our analysis revealed a gradual displacement of CTD from the CCD_Y99H/A128T_ + PIR complex compared to the WT CCD-PIR-CTD complex (Fig. 8A, Fig. S6, Movie S1), the CTD displacement was significantly reduced in the context of the CCD_Y99H/A128T_-EKC110-CTD complex compared to its WT CCD-EKC110-CTD counterpart (Fig. 8B, Fig. S6). Although, the CTD was slightly displaced from CCD_Y99H/A128T_-EKC110-CTD throughout the simulation compared to the WT CCD-EKC110-CTD complex, these shifts in orientation were transient and the CTD domain returned to its WT-like orientation in the presence of EKC110 but not in the presence of PIR (Fig. S6).

To unambiguously characterize the CTD displacement, we measured the internal volume of the CCD-CTD binding pocket throughout the simulations (Fig. S7). For the Y99H/A128T IN in complex with PIR, the CCD-CTD binding pocket exhibited an initial volume of 375 Å^3^, however, by the end of the simulation the volume increased to 433 Å^3^. This volume increase is correlated to the displacement of the CTD domain from the CCD_Y99H/A128T_-PIR-CTD complex compared to the WT CCD-PIR-CTD structure. Throughout 1 μs MD simulations, the WT CCD-PIR-CTD complex exhibited an average interface volume of 377.05 Å^3^ with an uncertainty of 49.30 Å^3^, while the average volume of the CCD_Y99H/A128T_-PIR-CTD interface increased to 396.87 Å^3^ with a uncertainty of 54.25 Å^3^. A Student T-test comparisons of the volume distributions yielded a p-value of 3.79 × 10^−211^ indicating that there is a statistically significant difference between the two volumes.

The CCD_Y99H/A128T_-EKC110-CTD complex exhibited an initial volume of 375 Å^3^ and after 1 μs MD simulations, the volume was largely unchanged (372 Å^3^), which is consistent with the more stable nature of the CCD-EKC110-CTD interface (32). For the CCD_Y99H/A128T_-EKC110-CTD system, we measured an average volume of 372 Å^3^ and a standard deviation of 58.48Å^3^; which were reduced from the average volume of 391.36 Å^3^ and standard deviation of 54.17Å^3^ measured from the WT CCD-EKC110-CTD complex. The Student T-test revealed a p-value of 1.42 × 10^−119^ indicating a statistically significant difference between the volume distributions. This reduction in volume can be attributed to the Y99H substitution, as replacing a Tyr with the less bulky His allows α-helix 1 from one CCD subunit and α-helix 5 from the partner CCD subunit to be packed closer together.

Taken together, the MD simulation results (Fig. 8, Figs. S6, S7) extend our experimental findings by indicating that Y99H/A128T mutations lead to changes in observed volumes of the ALLINI binding pocket (6)at the CCD-ALLINI-CTD interface and more readily displace CTD from the CCD_Y99H/A128T_ + PIR than the CCD_Y99H/A128T_ + EKC110 complex.

### FEP calculations

To quantify how Y99H, A128T and Y99H/A128T mutations affected CCD-ALLINI-CTD interactions we performed FEP calculations (Fig. S8). For this, we computed the relative free energy difference (ΔΔG) between the WT CCD vs CCD containing Y99H, A128T and Y99H/A128T mutations for their ability to form the CCD-ALLINI-CTD complexes (Fig S8). The single Y99H mutation carried a similar energetic penalty for both CCD-PIR-CTD (ΔΔ*G* = 0.65 ± 0.07 *kcal*/*mol*) and CCD-EKC110-CTD (ΔΔ*G* = 0.96 ± 0.86 *kcal*/*mol*) complexes, whereas the single A128T mutation yielded a much higher ΔΔG for the CCD-PIR-CTD complex (ΔΔ*G* = 5.24 ± 0.47 *kcal*/*mol*) compared to the free energy differences seen for the CCD-EKC-CTD complex (ΔΔ*G* = 2.09 ± 0.18 *kcal*/*mol*). The drug resistant mutations induced a free energy change of ΔΔ*G* = 7.34 ± 1.01 *kcal*/*mol* for the CCD_Y99H/A128T_-PIR-CTD complex, which was higher than the free energy change of ΔΔ*G* = 5.07 ± 0.74 *kcal*/*mol* measured for the CCD_Y99H/A128T_-EKC110-CTD complex. These findings suggest that the Y99H/A128T mutations are more unfavorable for PIR than EKC110.

## DISCUSSION

Our multidisciplinary studies have elucidated an unexpected mechanism of the viral resistance to PIR. Even though both Tyr99 and Ala128 are located within the V-shaped cavity at the CCD dimer, the Y99H/A128T mutations did not substantially affect direct binding of PIR to the CCD dimer or functional oligomerization of the full-length IN. Instead, these drug-resistant mutations introduced steric hindrance at the PIR mediated CCD-CTD interface and impaired the ability of the CCD_Y99H/A128T_ + PIR complex to bind CTD. Consequently, full-length IN_Y99H/A128T_ was substantially more resistant to the PIR induced hyper-multimerization than its WT counterpart. PIR was >150-fold less potent against HIV-1_(Y99H/A128T IN)_ vs the WT virus.

Cell culture based viral breakthrough assays with different ALLINIs consistently identified various drug resistance mutations in the vicinity of the inhibitors’ binding site on CCD (1, 3, 5, 7, 8, 14, 29, 34, 35). By contrast no mutations were detected within CTD. These findings suggest that HIV-1 is more tolerant to the drug-resistant mutations within the V-shaped cavity at the CCD dimer interface than at the complementary CTD interface which is composed of the invariant residues (32). Indeed, the Y99H/A128T IN changes only partly (∼2-fold) reduced HIV-1 infectivity (Fig. 1B), whereas the mutations of the key CTD residues that engage with the CCD-ALLINI complex are detrimental for the virus (36, 37).

The A128T change is the most frequently detected resistance mutation against different ALLINI chemotypes (1, 5, 6, 14, 34). Previous mechanistic studies with this IN mutation helped to delineate that the primary mode of action of ALLINIs was through inducing hyper-multimerization of IN rather than inhibiting IN binding to LEDGF/p75 (14). Indeed, the A128T change did not detectably affect the ALLINI IC_50_ values for IN-LEDGF/p75 binding. Instead, IN_A128T_ was substantially more resistant to the inhibitor induced hyper-multimerization than WT IN (14). However, the previous structural studies were limited to the ALLINI-CCD interactions and the underlying mechanism for the A128T IN resistance remained obscure (14). Our studies here reveal the importance of the ALLINI induced CCD-CTD interface for the emergence of the Y99H/A128T IN resistant viral phenotype. In turn, these findings raise a possibility that a number of previously reported resistant mutations that arise in response to different ALLINI chemotypes could also affect the inhibitor induced CCD-CTD interactions. In this regard, our biochemical assay reported here (Fig. 4) could offer a robust tool to examine the mechanisms of other drug-resistance mutations with respective ALLINIs.

Our structural and mechanistic studies with the viral resistance to PIR provided us with a means to rationally modify the parental compound to develop its improved analog EKC110. The removal of 3-methyl group from the pyrrolopyridine ring of PIR allowed EKC110 to reposition deeper inside the V-shaped cavity at the CCD dimer interface. In addition, our crystal structures (Fig. 7B) revealed a considerable repositioning of the CTD between EKC110 and PIR co-crystal structures, which could afford more space for the former complex to minimize steric clashes at the CCD-CTD interface induced by the drug-resistant mutations. Accordingly, EKC110 was more potent against IN_Y99H/A128T_ *in vitro* and HIV-1_(Y99H/A128T IN)_ in infected cells compared to the parental PIR. These exciting results inform future efforts to develop second generation ALLINIs with an enhanced barrier for resistance for their potential clinical use.

## MATERIALS AND METHODS

### Cell lines, virus infectivity and antiviral assays

HEK293T (ATCC) and HeLa TZM-bl (NIH AIDS Reference and Reagent Program) cells were cultured in Dulbecco’s modified eagle medium (DMEM, Gibco) supplemented with 10% fetal bovine serum (FBS, Sigma–Aldrich) and 1% penicillin–streptomycin (PS, Gibco). Cells were maintained in incubator at 37 °C and 5% CO_2_. Cell lines used in this study were tested monthly for Mycoplasma contamination.

For virus infectivity assay, HEK293T cells (2-4 x 105 cells/well in 6-well plate) were seeded one day prior to transfection of 2 µg replication competent pNL4-3 plasmid containing WT or mutant INs using HilyMax transfection reagent (Dojindo Molecular Technologies, Inc.) in 1:3 ratio. The medium was replaced with fresh medium at 12-16 h post-transfection and incubated at 37 °C. Then, 48 h post-transfection virus containing supernatant were collected, clarified, and filtered through 0.45 µm filter and the level of p24 was quantified by western blot. We used p24 normalized filtered viral supernatant from 293T cells to infect TZM-bl cells (seeded at 50000 cells/well in 24-well plate), incubated at 37 °C for 3-4 h, the medium was removed and replaced with fresh medium. The cells were collected at 48 h post-infection and virus infectivity were measured by luciferase assay (Promega).

For antiviral assay, full replication cycle experiments were performed with HIV-1_NL4.3_ containing WT or mutant INs as described (3, 38). Briefly, viruses were prepared in the presence of PIR or EKC110 or DMSO as a control. Target cells were pre-incubated with PIR or EKC110 or DMSO as a control, infected with viruses for 3-4 h at 37 °C, medium was replaced, and fresh inhibitors added. 48 h post infection, cells were collected, and infectivity were measured by luciferase assay. Effective concentration (EC_50_) of the inhibitors were calculated using Origin software (OriginLab, Inc.). HEK293T and HeLa TZM-bl cells were used as producer and target cells, respectively. All virus infections were performed in the presence of 8 µg/ml polybrene, and values were expressed as mean ± standard deviation (SD).

### Synthesis of ALLINIs

PIR was synthesized as described (5). EKC110 (compound **19)** was prepared following the synthetic procedure outlined in Scheme **1** (see Supplemental materials). Intermediate **2** was obtained by reacting commercially available diethyl malonate (**1**) and acetonitrile in presence of tin(IV) chloride (SnCl_4_) (39), while intermediate **6** was obtained by bromination of (1-methyl-1*H*-pyrazole-4-yl) methanol (**3**) (40) in presence of 33% HBr in acetic acid followed by reaction with 5-methyl-2-pyrrolidinone (**5**) in presence of NaH. Coupling of compounds **2** and **6** in presence of POCl_3_ and subsequent cyclization in presence of NaOEt afforded compound **8** (41). Oxidation of compound **8** with DDQ in toluene gave aromatized product **9** which was converted to the corresponding triflate **10** by reaction with triflic anhydride (Tf_2_O) in presence of triethylamine. Subsequent palladium mediated Suzuki coupling with 4-chlorophenylboronic acid in the presence of potassium carbonate and Pd(PPh_3_)_4_ gave intermediate **11**. Aldehyde derivative **13** was obtained by first, reduction of the ethyl ester to the corresponding alcohol with DIBAL-H followed by oxidation with pyridinium chlorochromate. Reaction of aldehyde **13** with trimethylsilyl cyanide in presence of ZnI_2_ gave silylated cyanohydrin **14** which, after hydrolysis with H_2_SO_4_ in methanol, provided hydroxyester **15**. Oxidation of the hydroxyl group with DMP followed by asymmetric reduction, using Corey-Bakshi-Shibata reagent ((*R*)-Me-CBS borane), gave the chiral alcohol **17**.^3^ Alkylation of **17** in presence of *t*-butyl acetate and perchloric acid and further saponification of intermediate **18** led to target compound **19 (EKC110)**.

### Recombinant proteins

Y99H/A128T mutations were introduced in the full-length IN, CCD and CTD-CCD constructs by PCR-directed mutagenesis and the proteins were expressed in BL21 (DE3). Full-length IN and CCD proteins were purified as described (11). The CTD-CCD proteins were prepared as described (32). Purified proteins were examined using NuPAGE Bis-Tris 4-12% acrylamide gels with MES as the running buffer (Invitrogen). Proteins were stained AcquaStain Protein Gel Stain (Bulldog-Bio).

### Analytical SEC

Recombinant WT and mutant IN proteins were analyzed using Superdex 200 10/300 GL column (GE Healthcare) with the running buffer containing 20 mM HEPES (pH 7.5), 1 M NaCl, 10% glycerol and 5 mM BME at 0.3 mL/min flow rate. The protein stocks were diluted to 20 µM IN with the running buffer and incubated for 1 h at 4 °C followed by centrifugation at 10,000 x g for 10 min. To estimate multimeric state of IN proteins we used the following standards: bovine thyroglobulin (670,000 Da), bovine gamma-globulin (158,000 Da), chicken ovalbumin (44,000 Da), horse myoglobin (17,000 Da) and vitamin B12 (1,350 Da). Retention volumes for different oligomeric forms of IN were as follows: tetramer ∼12.5 mL, dimer ∼14 mL, monomer ∼15-16 mL.

### SPR

The SPR biosensor binding experiments were performed using the Biacore^TM^ T200 (Cytiva). A nitrilotriacetic acid (NTA) sensor chip was conditioned with 350 mM NiSO_4_ at a flow rate of 30 µL/min for 1 min. His_6_-CCD and His_6_-CCD_Y99H/128T_ proteins containing C-terminal hexa-His-tag were immobilized on the NTA sensor chip to about 2,000 response units. The running buffer contained 0.01 M HEPES (pH 7.4), 0.15 M NaCl, 0.05% v/v Surfactant P20 (Cytiva), and 5% DMSO. The desired concentrations of inhibitors were prepared by serially diluting the compounds in 100% DMSO and then by adding the running buffer (without DMSO) to reach a final DMSO concentration of 5%. The sensor chip was regenerated with 350 mM EDTA. For each interaction, background binding and drift were subtracted via a NTA reference surface. Data were analyzed using Biacore T200 Evaluation software and fit with a 1:1 kinetic model. The sensorgrams were plotted using Origin software.

### DLS

The DLS assays were performed on a Malvern Zetasizer Nano s90 as described (3). Full length WT and mutant INs were analyzed at 200 nM in the presence of 500 nM PIR. Kinetic analysis was carried out at specified time points. In short, the reactions were performed in the DLS buffer (1 M NaCl, 2 mM MgCl_2_, 2 mM DTT, 50 mM HEPES, pH 7.5) which was filtered twice using 0.2 µm filter. Stock solution of PIR (1 mM) were prepared in filtered DMSO. 0.2 μL of PIR (1 mM) was added to 40 μL of IN (200 nM, diluted in DLS buffer) and size distributions of the mixture were recorded at 1, 5, 10, and 15 min. For a negative control, the same amount of IN was mixed with 0.2 µL filtered DMSO (100%).

### CTD binding to the CCD + ALLINI complex

2 µM His_6_-CCD and His_6_-CCD_Y99H/A128T_ proteins were immobilized on Ni-NTA resin in the binding buffer containing 50 mM HEPES (pH 7.5), 200 mM NaCl, 2 mM MgCl_2_, 35 mM imidazole, 0.1% (v/v) Nonidet P40 and 0.1% BSA. Subsequently, 2 µM CTD was added in the absence or presence of 2 µM PIR and the mixtures were rotated for 30 min using Tube Revolver Rotator at a speed of 40 rpm for 30 min. The resins were washed three times with the binding buffer to remove unbound proteins, and the bound proteins were separated by SDS–PAGE electrophoresis and visualized by staining with Coomassie-Blue-like AcquaStain (Bulldog-Bio).

### X-ray crystallography

The CCD and CCD_Y99H/A128T_ proteins were concentrated to 5 mg/mL and crystallized at 4 °C using the hanging drop vapor diffusion method as described previously (42). 2 μL protein was mixed with 2 μL reservoir, with 500 μL reservoir solution in the well which contained 0.1 M (NH_4_)_2_SO_4_, 0.1 M sodium cacodylate (pH = 6.5), 10% PEG 8000, and 5 mM DTT. The cubic-shaped crystals reached 0.1-0.2 mm after 1-2 weeks. The soaking buffer was prepared the same as the mother liquid but supplemented with 30% mixture of ethylene glycol, DMSO, and glycerol (1:1:1). The CCD and CCD_Y99H/A128T_ were soaked with either PIR or EKC110 (0.28 mM) in this cryoprotectant solution overnight before being flash-frozen in liquid nitrogen. Diffraction data were collected at 100 K by a Rigaku Micromax 007 with a Pilatus 200K 2D area detector at University of Colorado Anschutz Medical Campus X-Ray Crystallography Facility.

For the CTD-CCD + EKC110 crystal structure we used 10 nM stock of EKC110 in DMSO. To prepare the protein-drug complexes, The CTD-CCD construct contained solubilizing F185K/W243E IN mutations as described (32). The protein was diluted to 0.6 mg/mL by buffer containing 20 mM Tris-HCl pH 7.5, 0.5 M NaCl, and then supplemented with 25 µM EKC110 in the presence of 5% (v/v) glycerol. Following incubation on ice for 10 mins, the complexes were concentrated to 5 mg/mL using 10 kDa cutoff VivaSpin device (Satorius). The crystals grew at room temperature (23 °C) by adding 1 µL protein with 1 µL of reservoir containing 30 mM magnesium chloride, 30 mM calcium chloride, and 0.1 M imidazole-MES (Morpheus buffer system 1; Molecular Dimensions product code MD2-100-100), pH 6.5, 10% (w/v) PEG 8000, and 20% ethylene glycol. Crystals cryoprotected in the mother liquor supplemented with 30% glycerol were frozen by plunging them into liquid nitrogen.

### Structural studies

Data integration and reduction were performed with XDS (43). Molecular Replacement software Phaser (44) in the phenix (45) package was employed to solve all protein and ligand structures. Coot (46) and phenix.refine were used afterwards to refine structures. TLS (47) and restraint refinement was done for the last step of structure refinement.

The CCD + ALLINI crystals belonged to space group P3_1_21 with cell dimensions: a = b = 72.09 and c = 65.91 Å with a 18.84 KDa monomer in the asymmetric unit. The structures were refined to approximately 1.9-2.1 Å with R_work_ = 0.22 - 0.26 and R_free_ = 0.26 - 0.30. PDB entry 6NUJ was used as the starting model, and CCD structures in complexes with PIR and EKC110 were deposited on PDB with codes 8D3S and 8S9Q, respectively. CCD_Y99H/A128T_ complexed with PIR and EKC110 were deposited on PDB with codes 8T52 and 8T5A, respectively.

The CTD-CCD + EKC110 crystals belonged to space group P12_1_1 with cell dimensions: a = 61.954, b = 69.984, and c = 63.858 Å with a 51.74 KDa dimer in the asymmetric unit. The structure was refined to about 2.08 Å with R_work_ = 0.24 and R_free_ = 0.26. PDB entry 8A1Q of which CTD-CCD is complexed with PIR (32) was used as the starting model for the CTD-CCD + EKC110 structure (PDB with code 8T5B).

### MD simulations

As a starting point for all MD simulations, we utilized the crystal structure of PIR + WT CCD-CTD (PDBID: 8A1Q) and modeled the disordered chain regions not resolved in the CCD domain of the structure: residues 145 to 148 of CCD subunit 1 and residues 141 to 147 of CCD subunit 2, using Modeller (48). Subsequently, an all-atom model for Apo CCD dimers was derived by removing PIR from its complex with the inhibitor (PDBID: 8A1Q). The initial structure for the WT CCD dimer in complex with EKC110 was derived from the CCD-CTD + PIR complex by alchemically transforming the bound PIR molecules into EKC110 by substituting a methyl group from the pyrrolopyridine-based aromatic scaffold of PIR to a hydrogen. For all models, we added hydrogens to HIV-1 IN according to the protonation state of the amino acids at pH 7.0 as predicted by propKa (49), while maintaining the Mg^2+^ ions from the crystal structures. These models were then prepared for molecular simulation by solvating each system with TIP3P water molecules into a periodic box and ionized with Na^+^ and Cl^-^ ions to achieve a concentration of 150mM in VMD (50). The final simulation domains contained 142,889 and 142,883 atoms for the IN complexes with PIR and EKC respectively, with overall system dimensions of 115 Å x 108 Å x 119 Å.

In addition, for the 1 μs MD simulations of CCD_Y99H/A128T_-CTD complexed with PIR and EKC, we used the mutator plugin in VMD to introduce the mutations in the described structure (PDBID: 8A1Q), then, we derived coordinates for the ALLINIs into the CTD-CCD binding pocket in the same position as the WT structures and kept the Mg^2+^ ions. Structures for the CCD_Y99H/A128T_-CTD in complex with ALLINIs were then solvated and ionized following the same procedure described in the previous paragraph. The fully solvated models contained 144,082 atoms for the CCD_Y99H/A128T_-CTD in complex with PIR and EKC and system dimensions of 117 Å x 111 Å x 117 Å.

Prior to MD simulations, we performed the following equilibration procedure for all wild-type and Y99H/A128T IN complex systems (51). First, we energy minimized the solvent and ions around the protein while constraining the positions of protein and ligand atoms with a harmonic constant of 100 kcal/mol; the minimization procedure used the conjugate gradient scheme and was extended until the gradient converged to values below 10 kcal mol^-1^Å^-1^. Next, we thermalized the solvent and ions by slowly raising the temperature of the simulation domain from 50 K to 310 K at a rate of 0.5 K/ps while maintaining the constraints on the positions of protein and ligand atoms. A second energy minimization step followed, in which the restraints in the positions of protein and ligand atoms were released, allowing the positions of all atoms in the system to be optimized until the conjugate gradient converged to values below 10kcal mol^-1^ Å^-1^. This minimization procedure was followed by a second thermalization step where the positions of the protein backbone atoms were harmonically restrained with a light harmonic constant of 10 kcal/mol and the temperature of the simulation domain was slowly raised from 50 K to 310 K at a rate of 0.5 K/ps. Subsequently, we performed NPT equilibration simulations while the restraints on protein backbone atoms were slowly released at a rate of 2 kcal/mol/ns from 10 kcal/ to 0 kcal/mol over 5ns. For the equilibration simulations, we maintained the temperature at 310K using a Langevin thermostat with a thermal coupling constant of 1 ps^-1^ and a pressure of 1 atm via a Nose-Hoover barostat with a period of 100 ps and decay time of 50 ps.

After conducting the equilibration procedure, we performed 1 μs MD simulations in the NPT ensemble at a temperature of 310 K and pressure of 1 atm using the Langevin thermostat and Nose-Hoover barostat with the same parameters as above. Throughout all simulations we used a 2 fs timestep and periodic boundary conditions. Long range electrostatic interactions were calculated using the particle mesh Ewald method with a short-range cutoff of 12 Å and switching parameter of 10 Å. Throughout all MD and FEP simulations, the coordination number between the two Mg^2+^ ions and protein within 5Å in the CCD were constrained using the *coordNum* function in the Colvar module (52) of NAMD. All simulations were performed using the CHARMM36m force field parameters for proteins (53), the TIP3P model for water molecules (54). Force field parameters for both PIR and EKC were derived by analogy from the CHARMM general force field version 4.5 using CGenFF2.5 (55, 56). In total, summing the simulations for the wild-type and Y99H/A128T IN dimer systems in complex with PIR and EKC110 or in absence of ALLINIs, we compile a cumulative sampling of 5μs. All canonical MD simulations were performed in the NAMD3 molecular dynamics simulation engine taking advantage of GPU-accelerated computing (57).

### Binding pocket volume and orientation measurements

From the trajectories of 1 μs MD simulations for WT CCD-CTD and CCD_Y99H/A128T_-CTD in complexes with PIR and EKC110, we calculated the internal volume of the CCD-CTD binding pocket by defining as an outer shell of the protein atoms within 10 Å of the ALLINI bound and using *measure volinterior* plugin in VMD (58) for fuzzy-boundary volume detection with a grid spacing of 1Å, isovalue of 0.8, resolution of 5.5 and 64 rays casted by every voxel. Volumes reported are calculated using the 90-th percentile confidence threshold.

To quantify the displacement of the CTD domain from the CCD_Y99H/A128T_-PIR-CTD and CCD_Y99H/A128T_-EKC110-CTD complexes simulations with respect with their WT CCD-ALLINI-CTD counterparts, we measured the distance between the centers of mass of the CTD domains in the mutant and WT complexes after 1 μs molecular sampling by using the measure center command in VMD (50) and using the molecular mass of the atoms as weight. In addition, we used the package orient to calculate the principal axis of inertia of the CTD domains in the WT and mutant complexes and computed the angles of rotation between the axes in both complexes. We denote θ, Φ and Ψ, as the angles of rotation between the first, second and third principal axis of the CTD domains of the wild-type and mutant complexes throughout the text.

### FEP calculations

Alchemical FEP calculations were applied to the Y99H/A128T IN resistance mutations to quantify their effect on the binding of PIR and EKC110. Starting from the CCD_Y99H/A128T_-ALLINI-CTD and WT CCD + ALLINI complex crystal structures obtained in the present work, a dual-topology structure including the WT and mutant residues was created using the mutator plugin in VMD (50). These structures were then prepared for molecular simulation by solvating them in a TIP3P water box and ionizing them with Na^+^ and Cl^-^ ions to a salt concentration of 150 mM. Furthermore, all systems were subjected to the same equilibration procedure as the 1 μs MD simulations, described above, followed by a 15 ns post-restraint release equilibration step in the NVT ensemble at a temperature of 310K maintained via a Langevin thermostat with coupling constant of 1 ps^-1^. All FEP calculations were performed in the NVT ensemble with a temperature of 310 K and a Langevin thermostat coupling constant of 1 ps^-1^. All other simulation specifications and force field parameters were kept the same as in the long-scale MD simulations. All FEP calculations were performed using the NAMD2.14 molecular dynamics simulation engine (59).

The relative free energy differences were calculated using a thermodynamic cycle (Fig. S8B), where the vertical arms yield the free energy difference corresponding to the binding of PIR or EKC110 to WT or mutant IN (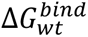 and 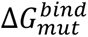), while the horizontal arms yield the free energy difference due to the residue substitution in the unbound HIV-1 IN and ALLINI-bound HIV-1 IN states (Δ*G^free^* and Δ*G^comp^*). In this manner, the relative free energy can be computed as

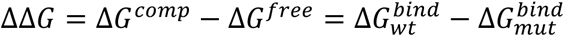

Here, we determined the free energy differences corresponding to the horizontal arms of the thermodynamic cycle (Δ*G^free^* and Δ*G^comp^*) via alchemical transformation of the residues using a dual-topology paradigm (60) in molecular dynamics simulations. In the dual-topology paradigm, we use a hybrid energy function:

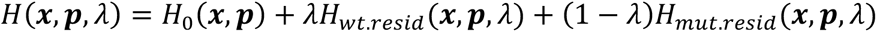

in which, *λ* is a coupling parameter connecting the physical wild-type (*λ* = 1) and mutant (*λ* = 0) states through alchemical states (0 < *λ* < 1). The FEP simulations were performed in a bidirectional approach by running 20 sequential equally spaced *λ*-windows in the forward direction from the WT to mutant IN followed by a simulation in the backward direction, from the mutant to WT IN (Fig. S8A). Each window of the alchemical transformation encompassed 1 ns of simulation, of which, 0.2 ns were used to equilibrate the simulation domain while the following 0.8 ns of sampling were used for the free energy calculations.

The free energy differences due to the residue substitution in the ALLINI-unbound IN system (Δ**G*^free^*) and in the ALLINI-bound IN system (Δ**G*^comp^*) were calculated from the forward and backward trajectories using the Bennet acceptance ratio estimator (61) as implemented in the ParseFEP plugin (62) in VMD (50). All FEP simulations were repeated in three independent replicates, the relative free energy differences (ΔΔ*G*) reported are the result of averaging the calculated ΔΔ*G* for the three independent replicates and the error bars represent the standard deviation between independent measurements (Fig. S8C).

### Data availability

The data presented in this manuscript are available from the corresponding authors upon reasonable request. The refined models and the associated X-ray diffraction data are deposited into the Protein Data Bank under accession codes 8S9Q (PIR + CCD), 8T5A (PIR + CCD_Y99H/A128T_), 8D3S (EKC110 + CCD), 8T52 (EKC110 + CCD_Y99H/A128T_) and 8T5B (CTD-CCD + EKC110).

## ACKNOWLEDGEMENTS

We are grateful to Dr. John Hardin at Structural Biology and Biophysics Core Facilities, University of Colorado Anschutz Medical Campus, and Dr. Jay Nix at ALS Beamline 4.2.2 for their support in collecting crystal diffraction data. We thank Dr. Daniel Adu-Ampratwum at the Ohio State University for his helpful advice throughout these studies and critical reading of the manuscript. This work was supported by NIH grants R01 AI143649 (to M.K. and J.R.F.), U54 AI170855 (to M.K.), U54 AI170791 (to J.R.P. and P.C.), AI 141327 (to B.K.), MH-116695 (to R.F.S.). This work was also funded in part by the Emory University Center for AIDS Research NIH grant P30-AI050409. The work in P.C. laboratory was also supported by the Francis Crick Institute, which receives its core funding from Cancer Research UK (CC2058), the UK Medical Research Council (CC2058), and the Wellcome Trust (CC2058). We acknowledge computational support through the Delaware Advanced Research Workforce and Innovation Network (DARWIN) as well as the Caviness cluster. This work used Stampede2 at TACC through allocation MCB-170096 from the Advanced Cyberinfrastructure Coordination Ecosystem: Services & Support (ACCESS) program, which is supported by National Science Foundation awards #2138259, #2138286, #2138307, #2137603, and #2138296. Support from the University of Delaware CBCB Bioinformatics Data Science Core Facility (RRID:SCR_017696) including use of the BioStore computational resources was made possible through funding from Delaware INBRE (P20GM103446), NIH Shared Instrumentation Grant (S10OD028725) the State of Delaware, and the Delaware Biotechnology Institute.”

