## Supplemental Materials for "The structural and mechanistic bases for the viral resistance to allosteric HIV-1 integrase inhibitor pirmitegravir"

[#] Contributed equally.

**Table S1. Data collection for the crystal structures**

| complexes | WT CCD + PIR | $CCD_{Y99H/A128T}$ + PIR | WT CCD + EKC110 | $CCD_{Y99H/A128T}$ + EKC110 | CTD-CCD + EKC110 |
|---|---|---|---|---|---|
| **PDB ID** | **8S9Q** | **8T5A** | **8D3S** | **8T52** | **8T5B** |
| X-Ray source | Rigaku Micromax 007 | Rigaku Micromax 007 | Rigaku Micromax 007 | Rigaku Micromax 007 | ALS 4.2.2 |
| Software | XDS | XDS | XDS | XDS | XDS |
| Wavelength | 1.54178 | 1.54178 | 1.54178 | 1.54178 | 1.00003 |
| Space group | P $3_1$ 2 1 | P $3_1$ 2 1 | P $3_1$ 2 1 | P $3_1$ 2 1 | P 1 $2_1$ 1 |
| Unit cell dimension<br>a, b, c (Å)<br>a, b, g (Å) | 72.0, 72.0, 65.9<br>90, 90, 120 | 71.8, 71.8, 66.0<br>90, 90, 120 | 72.5, 72.5, 66.2<br>90, 90, 120 | 72.2, 72.2, 66.4<br>90, 90, 120 | 62.0, 70.0, 63.9<br>90, 100.732, 90 |
| Resolution (Å) | 31.17 - 2.26<br>(2.341 - 2.26) | 31.56 - 1.933<br>(2.003 - 1.933) | 29.28 - 1.855<br>(1.921 - 1.855) | 31.28 - 2.075<br>(2.149 - 2.075) | 46.72 - 2.08<br>(2.154 - 2.08) |
| No. total reflection | 224392 (13449) | 21086 (78) | 29323 (456) | 21042 (356) | 59212 (4594) |
| No. unique reflection | 9576 (941) | 10545 (39) | 14677 (240) | 11041 (284) | 31574 (2807) |
| $R_{merge}$ | 0.288 (1.947) | 0.08091 (0.3051) | 0.02835 (0.2928) | 0.02927 (0.5911) | 0.06936 (0.9956) |
| $R_{pim}$ | 0.05948 (0.5318) | 0.08091 (0.3051) | 0.02835 (0.2928) | 0.02927 (0.5911) | 0.06936 (0.9956) |
| CC1/2 | 0.997 (0.506) | 0.993 (0.788) | 0.999 (0.913) | 0.999 (0.365) | 0.997 (0.32) |
| I/sI | 8.67 (0.82) | 7.97 (1.64) | 18.04 (1.87) | 28.93 (0.98) | 6.22 (0.52) |
| Completeness | 99.94 (100.00) | 69.77 (2.63) | 83.52 (13.78) | 87.16 (22.95) | 97.17 (84.72) |
| Multiplicity | 23.4 (14.3) | 2.0 (2.0) | 2.0 (1.9) | 1.9 (1.3) | 1.9 (1.6) |

Data were collected from a single crystal. Value in parentheses are from the highest-resolution shell

### Table S2: Refinement statistics of the crystal structures

| complexes | WT CCD + PIR | $CCD_{Y99H/A128T}$ + PIR | WT CCD + EKC110 | $CCD_{Y99H/A128T}$ + EKC110 | CTD-CCD + EKC110 |
|---|---|---|---|---|---|
| PDB ID | 8S9Q | 8T5A | 8D3S | 8T52 | 8T5B |
| No. reflection used in refinement | 9575 (941) | 10544 (39) | 14556 (235) | 10880 (280) | 31455 (2733) |
| No. reflection used for $R_{free}$ | 946 (95) | 568 (2) | 1449 (27) | 1104 (28) | 1998 (165) |
| $R_{work}$ (%) | 0.2551 | 0.2567 (0.4078) | 0.2820 | 0.2607 | 0.2403 (0.3660) |
| $R_{free}$ (%) | 0.2930 | 0.2784 (0.8347) | 0.3028 | 0.3091 | 0.2643 (0.3986) |
| No. non-hydrogen atoms<br>Protein<br>Ligan/ion<br>Water | 1130<br>1028<br>65<br>67 | 1102<br>1028<br>65<br>39 | 1081<br>1040<br>62<br>7 | 1108<br>1028<br>62<br>46 | 3399<br>3247<br>126<br>82 |
| Wilson B-factor | 36.03 | 25.20 | 29.75 | 27.21 | 35.79 |
| Average B-factor<br>Protein<br>Ligan/ion<br>Water | 43.19<br>42.99<br>38.17<br>48.80 | 31.44<br>31.44<br>30.30<br>35.17 | 36.95<br>37.12<br>33.26<br>28.78 | 36.69<br>36.73<br>32.57<br>38.76 | 43.64<br>43.89<br>34.46<br>41.65 |
| R.m.s deviations<br>Bond length (Å)<br>Bond angle (°) | 0.002<br>0.35 | 0.011<br>0.87 | 0.011<br>1.22 | 0.001<br>0.35 | 0.211<br>3.40 |
| Ramachandran<br>Favored (%)<br>Allowed (%)<br>Outliers (%) | 98.41<br>1.59<br>0.0 | 98.41<br>1.59<br>0.0 | 98.41<br>1.59<br>0.00 | 98.41<br>1.59<br>0.0 | 95.99<br>3.26<br>0.75 |
| Rotamer outliers (%) | 0.00 | 0.00 | 1.80 | 0.00 | 1.78 |
| Clash score | 1.43 | 3.81 | 0.94 | 0.48 | 4.23 |

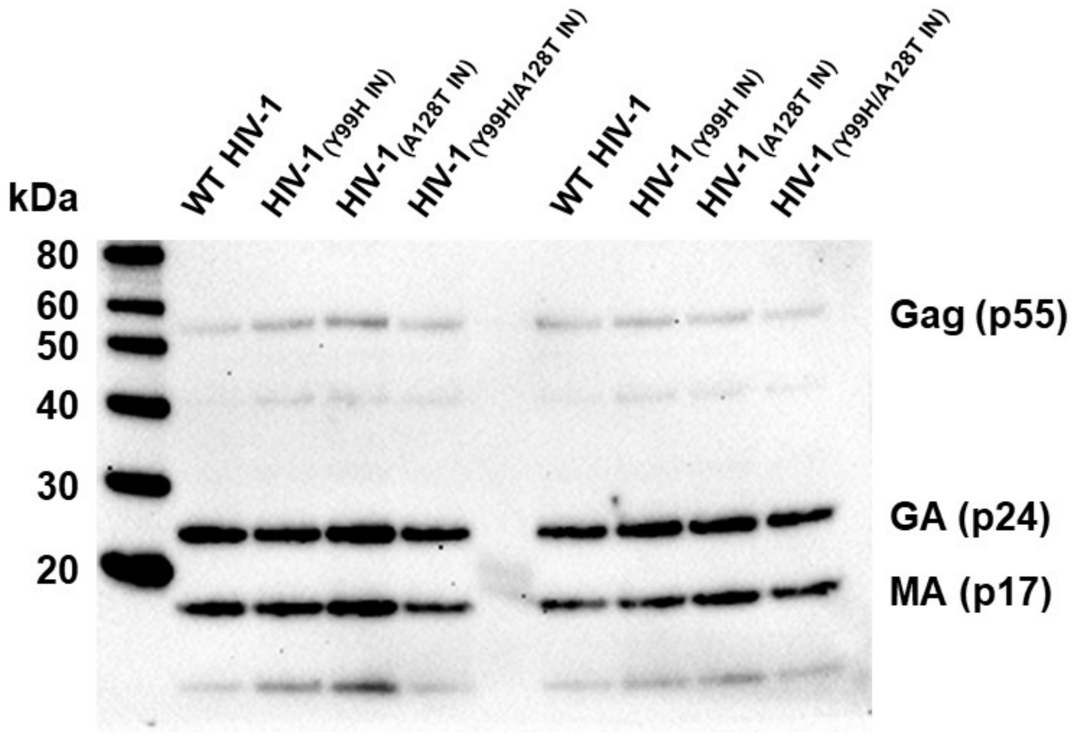


**FIG S1.** Virion productions for WT and indicated mutant viruses. Representative immunoblots showing capsid (p24) levels of WT and integrase mutant viruses. HIV-1 virions were produced in HEK293T cells by transfecting 2 μg of full-length pNL4-3 (WT or containing indicated mutations in IN). 48 h post-transfection viruses present in supernatant were collected, clarified, and filtered through 0.45 μm filter. The level of p24 in viral supernatant was analyzed by western blot using anti-HIV1 p55 + p24 + p17 antibody (ab63917) in ChemiDoc XRS+ System (bio-rad).

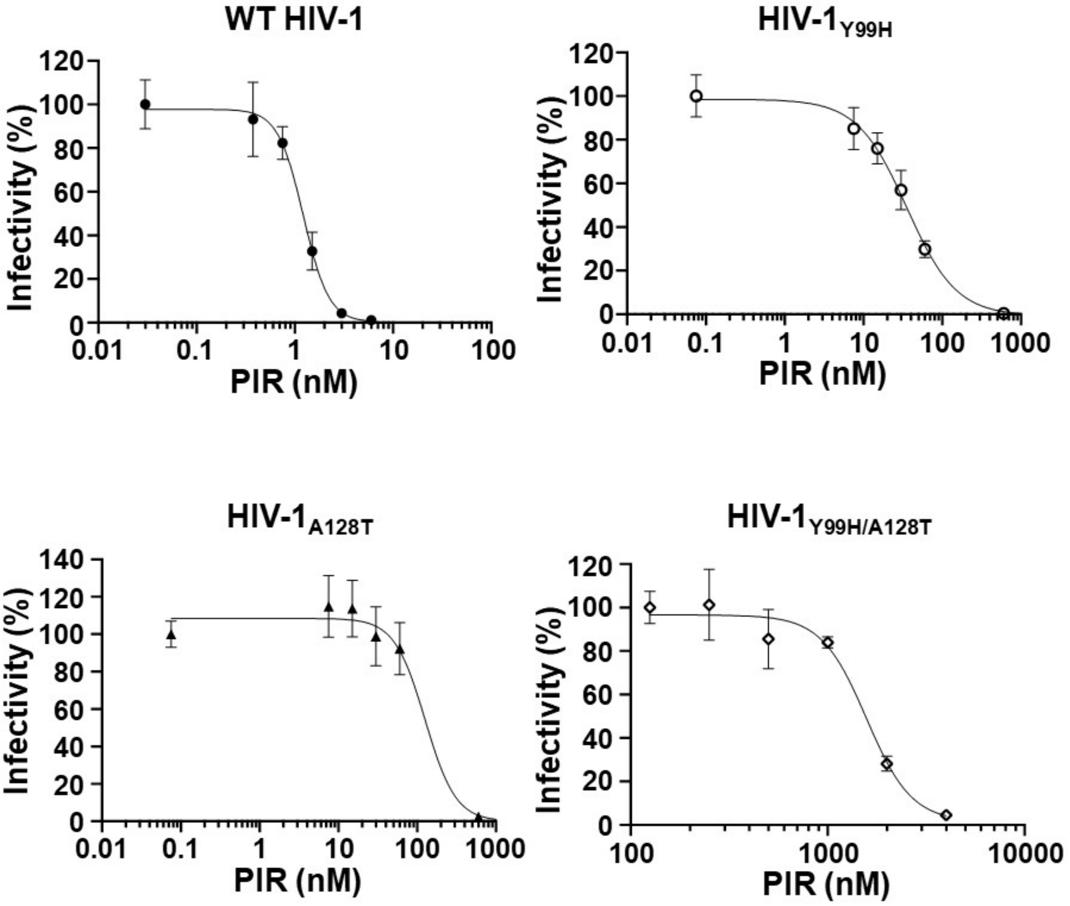


**FIG S2.** Antiviral activities of PIR against WT and IN mutant viruses. HIV-1 virions were prepared in the presence of PIR at indicated concentrations or DMSO as control in producer cells (HEK293T). Target cells (HeLa TZM-bl) were infected with pretreated virions in the presence of PIR at same concentrations or DMSO as control. 48 h post infection infectivity was measured by luciferase assay. Effective concentration ($EC_{50}$) of drugs were calculated using Origin software (OriginLab, Inc.) and are reported in Table 2.

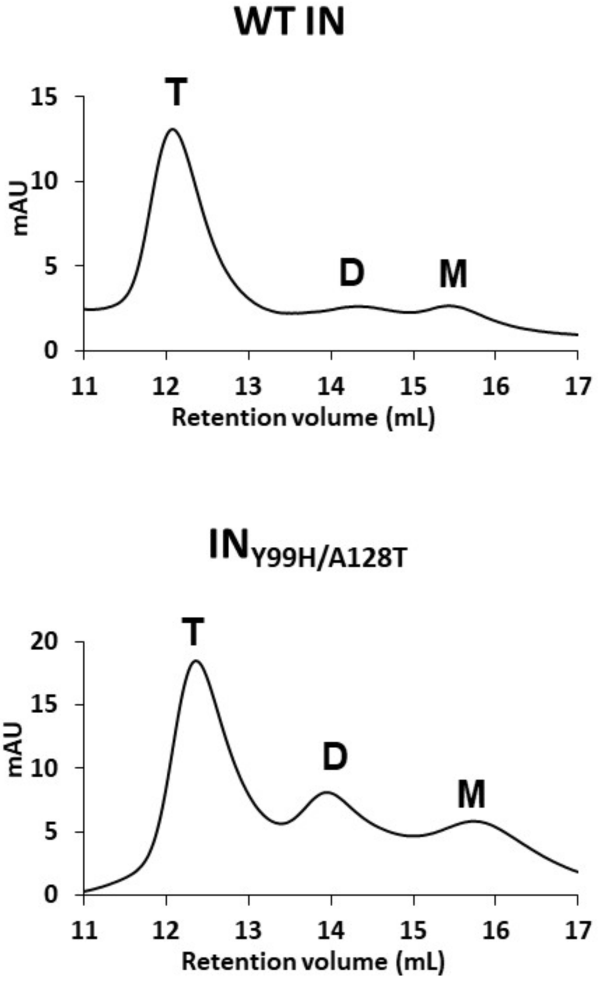


**FIG S3.** Analytical SEC of full-length WT IN (A) and $IN_{Y99H/A128T}$ (B). Elution chromatograms ($A_{280}$) of indicated recombinant HIV-1 IN from a Superdex-200 10/30 column are shown. The expected elution volumes for the tetrameric (T), dimeric (D), and monomeric (M) INs are indicated above each chromatogram.

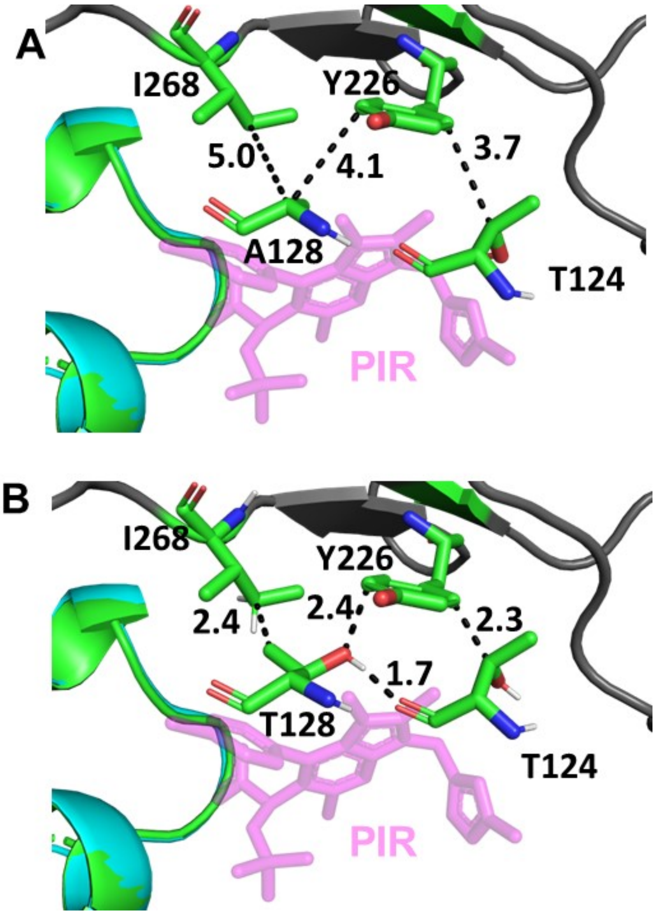


**FIG S4.** Analysis of PIR mediated CCD-CTD interfaces. (A) Distances between indicated CCD and CTD residues are shown in the structure of WT CTD-CCD + PIR to complement FIG 5B. (B) Distances between indicated CCD and CTD residues are shown when the structure of $CCD_{Y99H/A128T}$ + PIR is superimposed onto the structure of WT CTD-CCD + PIR to complement FIG 5C.

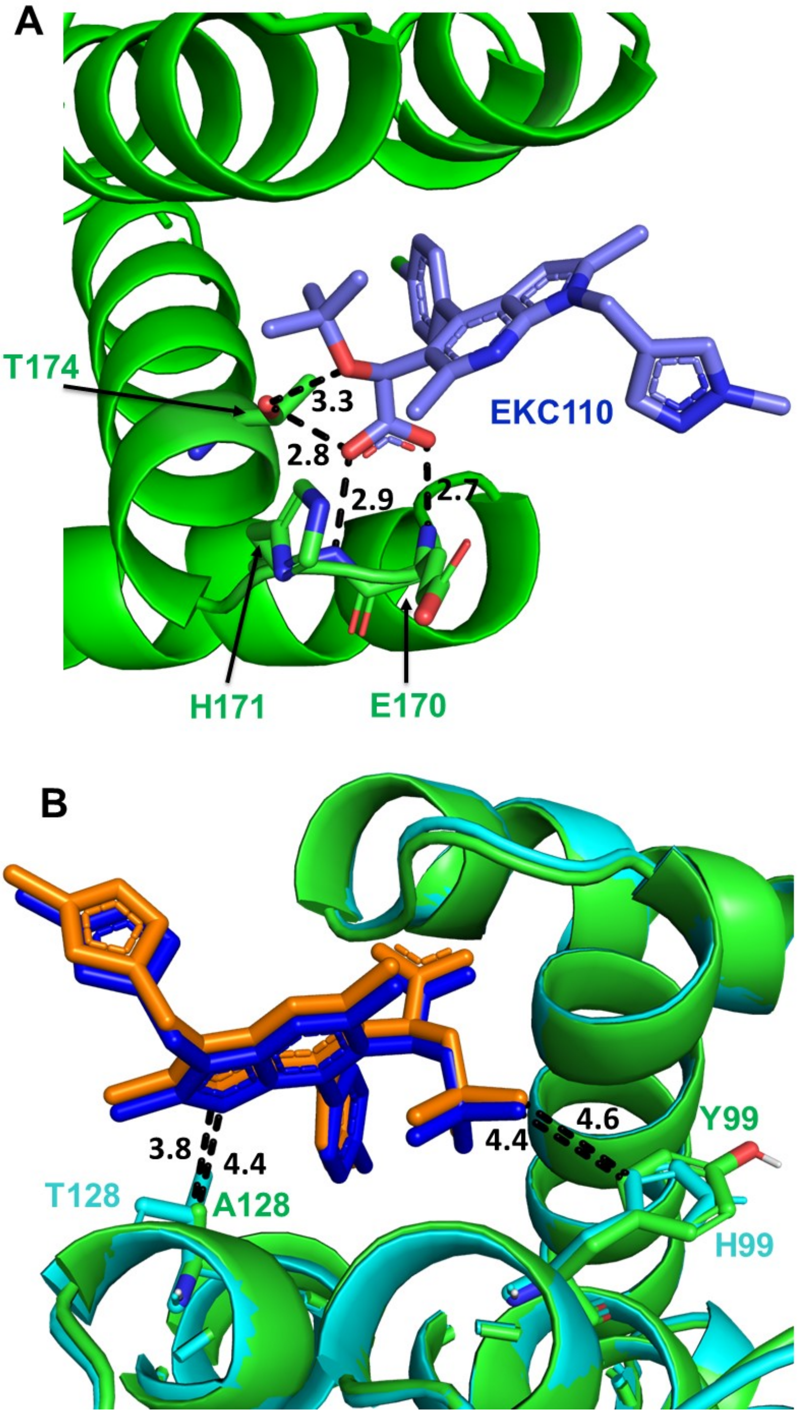


**FIG S5.** Interactions of EKC110 (blue) with the V-shaped pocket at the CCD dimer interface. (A) The crystal structure of EKC110 bound to WT CCD. Shown are bidentate hydrogen bonding established by the EKC pharmacophore carboxylate with backbone amides of Glu170 and His171. Furthermore, Thr174 side chain hydrogen bonds with the oxygen of the *tert*-butoxy moiety and carboxylate of EKC110. (B) comparative analysis of the crystal structures of EKC110 (blue) + WT CCD (green) vs EKC110 (orange) + $CCD_{Y99H/128T}$ (cyan). The closest distances between i) the EKC110 pyrrolopyridine ring to Cβ on either Ala128 or Thr128; ii) the EKC *tert*-butoxy moiety and the aromatic ring of Tyr99 or the imidazole of His99 are indicated.

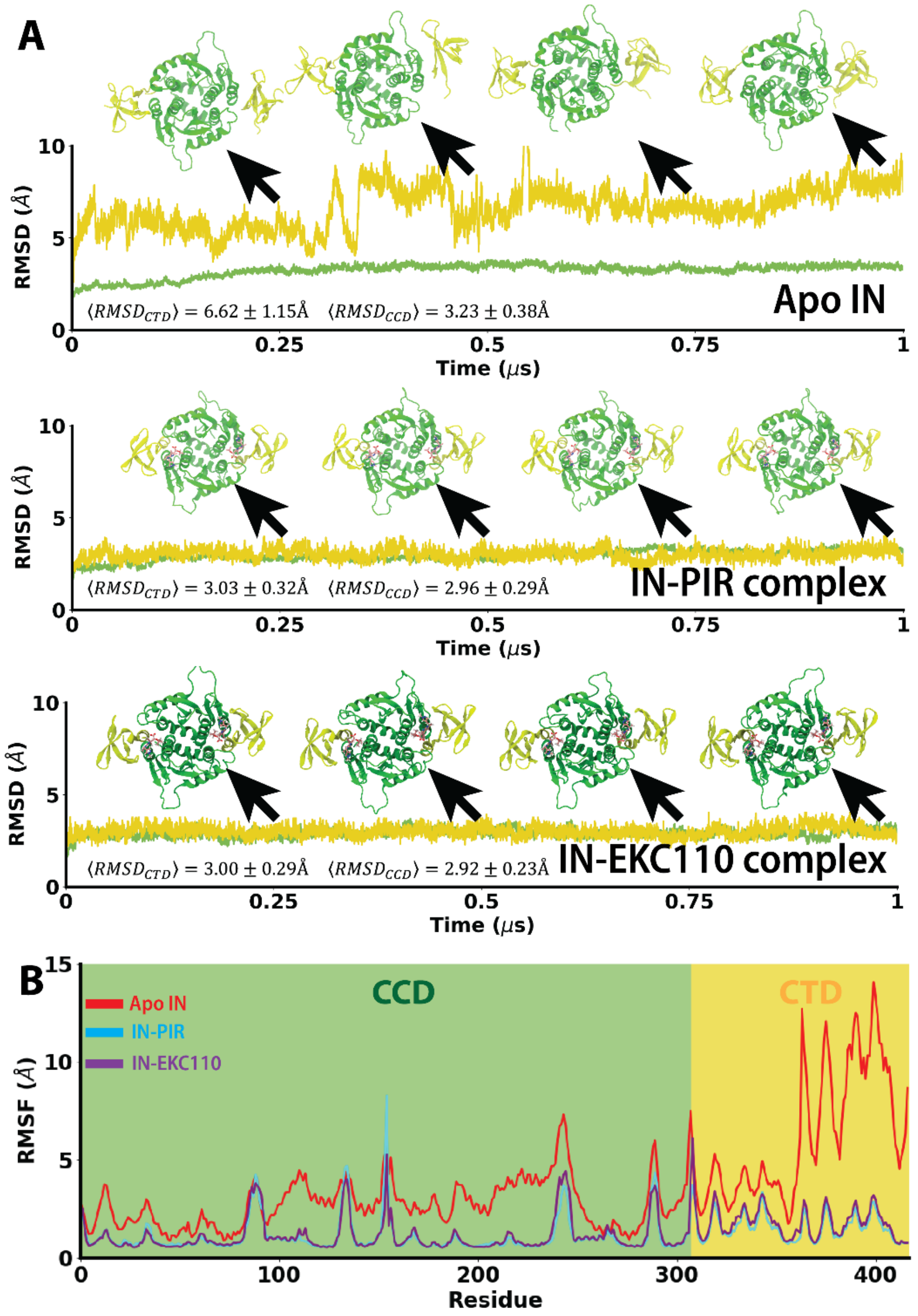


**FIG S6.** ALLINI effects on the CTD-CCD interactions. (A) Root means squared deviations (RMSD) calculated for the CCD (green) and CTD (yellow) domains of IN in Apo, PIR bound and EKC110 bound complexes though 1 μs MD simulation. B) Root mean squared fluctuation (RMSF) calculated for each residue in the absence of ALLINIs (red) or in the complex with PIR (cyan) or EKC110 (purple).

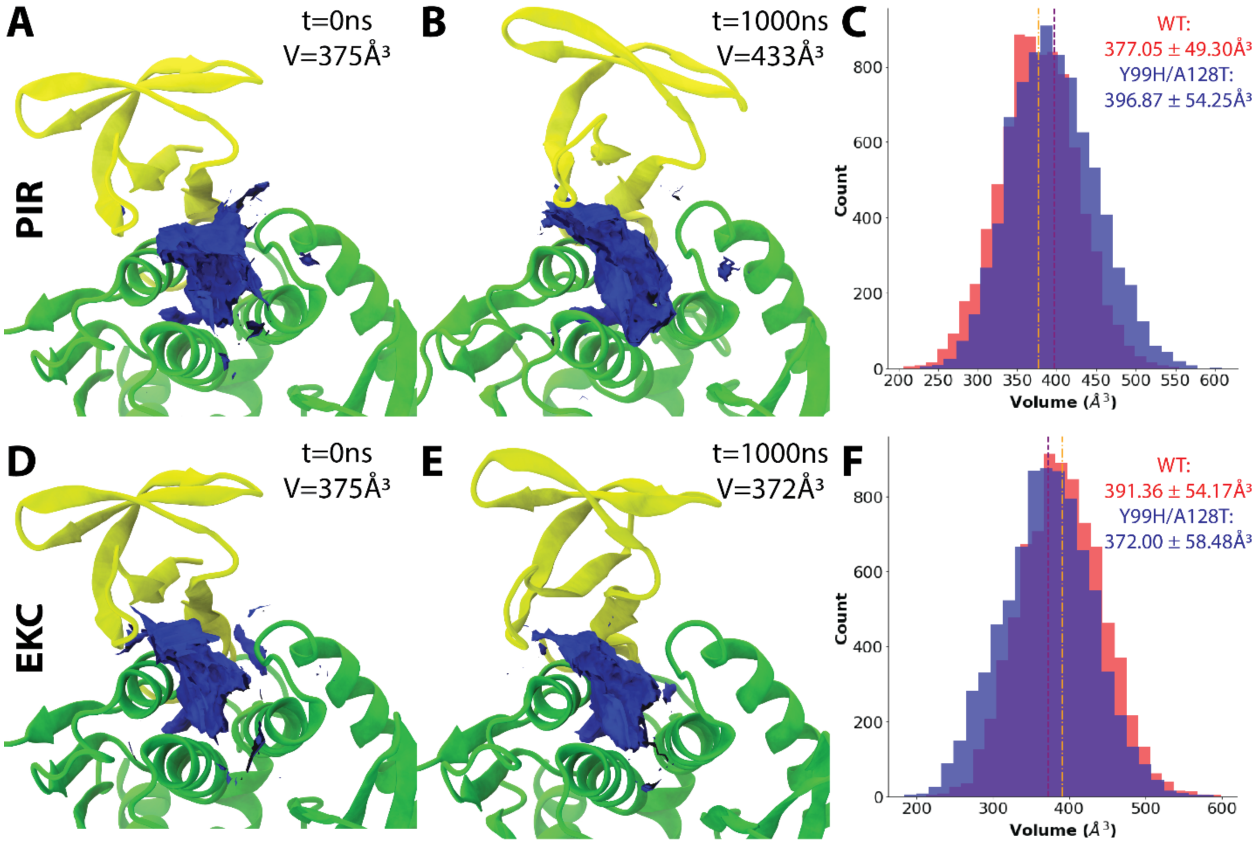


**FIG S7.** The CCD-ALLINI-CTD interface volume analyses. (A, D) Initial CCD-CTD interface volume for CCD(Y99H/A128T) in the complex with PIR (A) or EKC110 (D). (B, E) CCD-CTD interface internal volume after 1 μs simulation for CCD(Y99H/A128T) in the complex with PIR (B) or EKC (E). IN CCD and CTD domains are colored in green and yellow, respectively and the CCD-CTD interface volume in blue. (C, F) Internal volume distributions of the CCD-CTD interface for WT (blue) and the Y99H/A128T mutant (red) INs in complex with PIR (c) or EKC (f) over 1 μs of simulation.

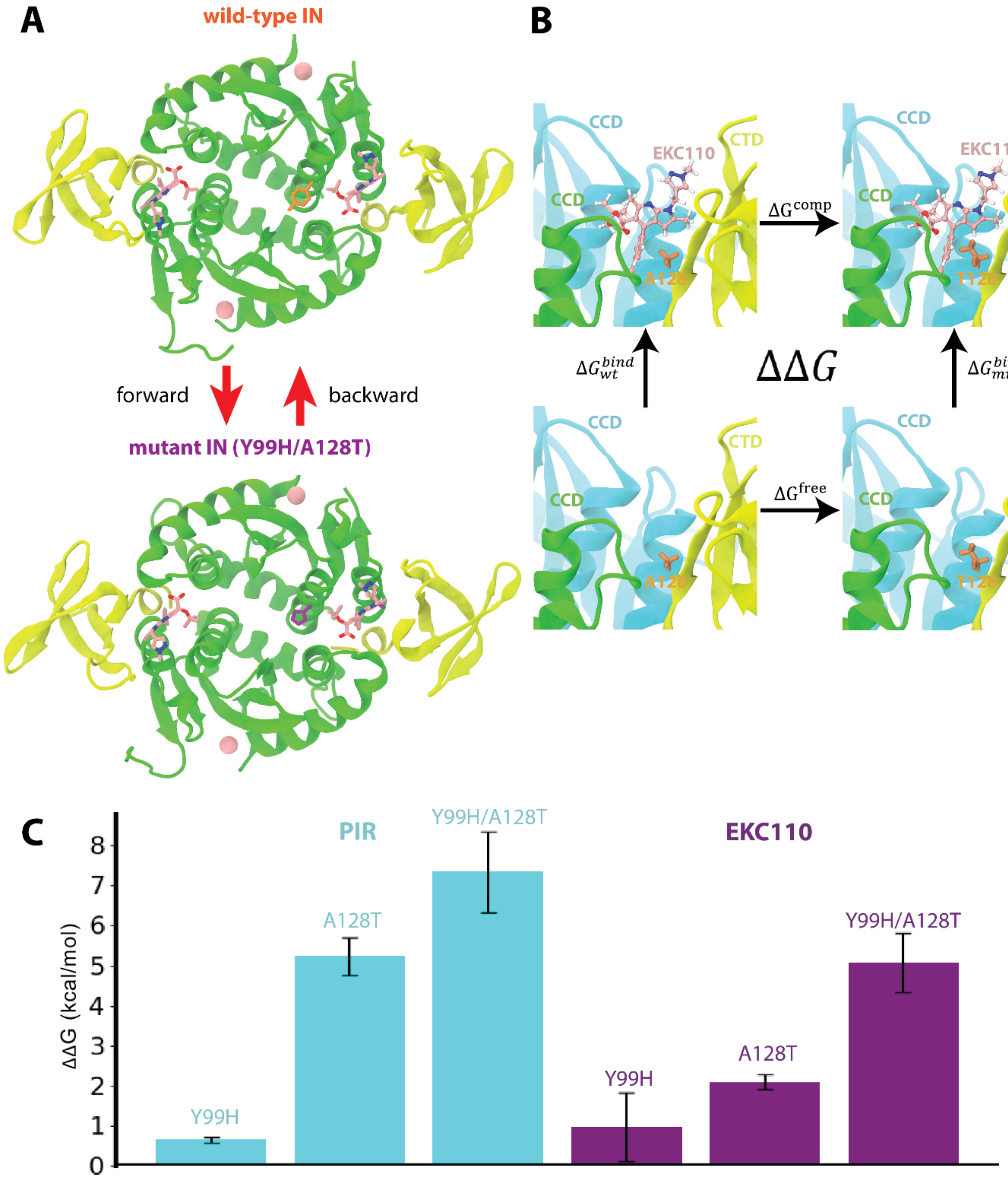


**FIG S8.** The free energy perturbation calculations. (A) The alchemical substitutions of WT IN residues to the drug resistant mutants were made in a forward (WT to mutant) and backward (mutant to WT) path. Snapshots for the initial and final states of the FEP trajectories for the Y99H/A128T IN mutations on the CTD-CCD + EKC110 are shown. CTD and CCD are colored yellow and green, respectively while the $Mg^{2+}$ is colored pink. WT Tyr99 and Ala128 residues are colored orange, while the mutated His99 and Thr128 are colored purple. (B) Thermodynamic cycle used to calculate relative free energy differences caused by CTD-CCD IN residue substitutions in complex with EKC110. The horizontal paths represent the alchemical residue substitutions, and the vertical paths represent the ALLINI binding. The example shows the alchemical mutation of E170A. IN CTD is colored in yellow and CCD subunits 1 and 2 are colored in green and cyan, respectively. (C) Relative free energy differences induced by the Y99H, A128T and Y99H/A128T mutations in the presence of PIR and EKC110 at the CTD-CCD interface.

**Scheme 1.** Synthesis of compound **19** (EKC110)

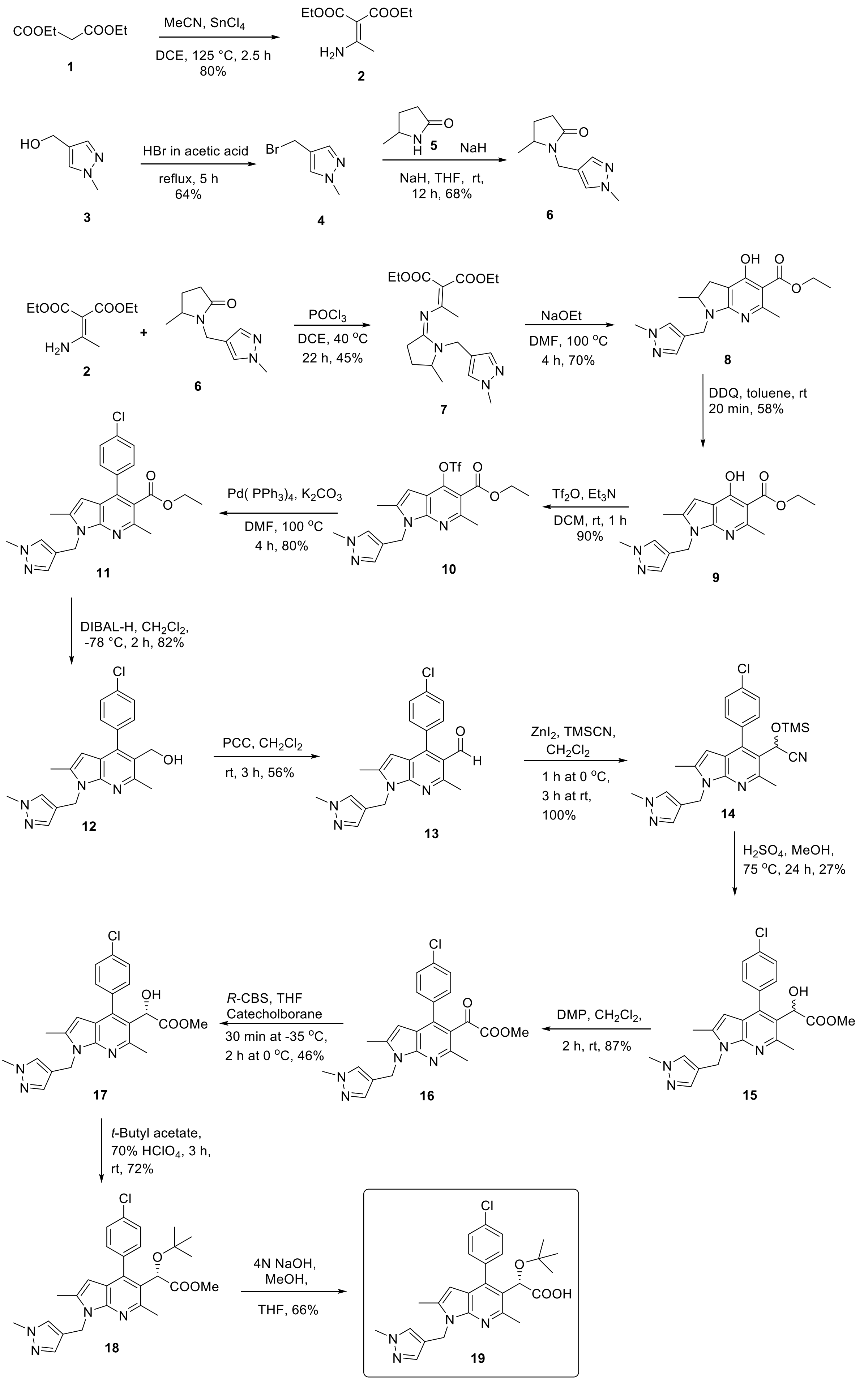

**Synthesis of compound 19 (EKC110)**

**General Procedures.** Anhydrous solvents were purchased from Aldrich Chemical Co., Inc. (Milwaukee, WI). All commercially available reagents were used without further purifications. Reagents were purchased from commercial sources. All the reactions were carried out under nitrogen in oven-dried glassware unless otherwise noted. Thin layer chromatography was performed on Analtech GHLF silica gel plates. Column chromatography was accomplished on Combiflash Rf200 or via reverse-phase high performance liquid chromatography. $^{1}H$, $^{13}C$, $^{19}F$, and $^{31}P$ NMR spectra were recorded on a Bruker Ascend 400 spectrometer at 25 °C (400 MHz, 101 MHz, 377 MHz and 162 MHz) as noted and residual proton solvent signals were used as internal standards. Deuterium exchange and decoupling experiments were utilized to confirm proton assignments. NMR processing was performed with MestReNova version 10.0.2-15465. Signal multiplicities are represented by s (singlet), d (doublet), dd (doublet of doublets), t (triplet), q (quadruplet), br (broad), bs (broad singlet), m (multiplet). Coupling constants (*J*) are in hertz (Hz). Mass spectra were determined on a Micromass Platform LC spectrometer using electrospray ionization. Purity of final compounds was determined to be >95%, using UPLC analyses performed on a Waters Acquity UPLC System with a Kinetex LC column (2.1 mm Å, 50 mm, 1.7 µm, C18, 100 Å) and further supported by clean NMR spectra. Mobile phase flow was 0.4 mL/min with a 1.20 min gradient from 95% aqueous media (0.05% formic acid) to 95% $CH_3CN$ (0.05% formic acid) and a 4.5 min total acquisition time. Photodiode array detection was from 190 to 360 nm.

**Diethyl 2-(1-aminoethylidene) malonate (2)**

To a solution of diethyl malonate **1** (19.05 mL, 124 mmol) and acetonitrile (6.46 mL, 124 mmol) in 1,2-dichloroethane (DCE) (80 mL) under nitrogen atmosphere was slowly added tin(IV) chloride ($SnCl_4$) (31.9 mL, 272 mmol) over 30 min and then the mixture was heated to 125 °C for 2.5 h. The mixture was concentrated to a paste and then dissolved in acetone (350 mL), transferred to a beaker, vigorously stirred and saturated $Na_2CO_3$/water (250 mL) was added dropwise to pH 9-10. The slush was filtered through a bed of celite, stirring the surface of the Celite® to facilitate filtering. The filter cake was washed with dichloromethane (4×200 mL). The aqueous phase was separated from the filtrate and the organic phase was dried ($Na_2SO_4$), concentrated and purified by column chromatography (90/10 to 80/20 hexane/ethyl acetate) to provide the desired compound **2** (20 g, 80%); $^1$H NMR (400 MHz, $CDCl_3$): $\delta$ 8.88 (s, 1H), 5.20 (s, 1H), 4.16 (dq, $J$ = 19.8, 7.1 Hz, 4H), 2.12 (s, 3H), 1.26 (dt, $J$ = 12.6, 7.1 Hz, 6H); $^{13}$C NMR (101 MHz, $CDCl_3$): $\delta$ 169.0, 168.6, 163.4, 93.2, 60.4, 59.7, 21.9, 14.4, 14.3; MS (EI, m/e) = 202 (M+1).

**4-(Bromomethyl)-1-methyl-1*H*-pyrazole (4)**

To a stirred solution of (1-methyl-1*H*-pyrazole-4-yl)methanol **3** (1.5 g, 13.37 mmol) in glacial acetic acid (7.5 ml) was added 33 % HBr in acetic acid (18 mL, 66.96 mmol) and the mixture was refluxed for 5 h. The solvent was removed under reduced pressure. The residue was crystallized from DCM and $Et_2O$ to afford the desired compound **4** (1.5 g, 64%); $^1$H NMR (400 MHz, $D_2O$) $\delta$ 8.00 (s, 2H), 4.50 (s, 2H), 4.00 (s, 3H); $^{13}$C NMR (101 MHz, $D_2O$) $\delta$ 135.1, 132.6, 122.8, 53.5, 37.9; MS (EI, m/e) = 198 ($M+Na^+$).

**5-Methyl-1-((1-methyl-1*H*-pyrazol-4-yl)methyl)pyrrolidin-2-one (6)**

To a solution of 5-methyl-2-pyrrolidinone **5** (755 mg, 7.61 mmol) in THF (30 mL) was added NaH (60 %, 610 mg, 15.22 mmol) at 0 °C. The mixture was stirred at rt for 30 min, then a solution of 4-(bromomethyl)-1-methyl-1*H*-pyrazole (**4**) (2 g, 11.41 mmol) in DMF (1 mL) was added. The resulting mixture was stirred at rt overnight. The reaction was quenched with $NH_4Cl$ (20 ml), diluted with ethyl acetate (20 mL) and then concentrated under vacuum. The residue was dissolved in ethyl acetate (50 mL) and washed with $H_2O$ (10 mL). The organic layer was dried over $MgSO_4$ and evaporated under reduced pressure. The crude residue was dissolved in acetonitrile (10 mL) and washed with hexane (10 mL x 2). The acetonitrile layer was concentrated and dried over $MgSO_4$ to afford the desired compound (**6**) (1 g, 68%); $^1$H NMR

(400 MHz, MeOD) $\delta$ 7.56 (s, 1H), 7.40 (s, 1H), 4.63 (d, $J$ = 15.2 Hz, 1H), 4.03 (d, $J$ = 15.2 Hz, 1H), 3.84 (s, 3H), 3.71 – 3.55 (m, 1H), 2.48 – 2.12 (m, 3H), 1.76 – 1.55 (m, 1H), 1.23 (d, $J$ = 6.3 Hz, 3H); $^{13}$C NMR (101 MHz, MeOD) $\delta$ 177.3, 139.8, 131.5, 118.2, 54.8, 38.8, 35.1, 31.2, 27.6, 19.9; MS (EI, m/e) = 194 (M+1).

**Diethyl (Z)-2-(1-((5-methyl-1-((1-methyl-1H-pyrazol-4-yl)methyl)pyrrolidin-2-ylidene)amino) ethylidene)malonate (7)**

To a solution of **6** (3.2g, 16.55 mmol) in 1,2-dichloroethane (DCE) (20 mL) under nitrogen atmosphere was added $POCl_3$ (2.32 mL, 24.82 mmol) dropwise over 15 minutes. The resulting mixture was stirred at ambient temperature for 1 h before addition of diethyl 2-(1-aminoethylidene)malonate (**2**) (4.33 g, 21.52 mmol). After heating the mixture at 40 °C for 22 h, saturated $NaHCO_3$/water was carefully added (20 mL) and the mixture was stirred at rt for 1 h then extracted with dichloromethane (3 x 40 mL). The combined organic phase was washed with brine, dried ($Na_2SO_4$), and concentrated under vacuum. The product was purified by flash chromatography using hexane/ethyl acetate (80/20 to 50/50) to afford the desired compound **7** (2.80 g, 45 %); $^1$H NMR (400 MHz, $CDCl_3$) $\delta$ 7.39 (s, 1H), 7.35 (s, 1H), 4.61 (d, $J$ = 15.1 Hz, 1H), 4.15 (q, $J$ = 7.0 Hz, 2H), 4.11 – 3.99 (m, 2H), 3.93 (d, $J$ = 15.2 Hz, 1H), 3.83 (s, 3H), 3.50 (q, $J$ = 7.0 Hz, 1H), 2.51 – 2.30 (m, 2H), 2.21 (s, 3H), 2.09 – 1.99 (m, 1H), 1.53 – 1.42 (m, 1H), 1.23 (t, $J$ = 7.0 Hz, 3H), 1.16 (ddd, $J$ = 15.1, 7.0, 1.2 Hz, 6H); $^{13}$C NMR (101 MHz, $CDCl_3$) $\delta$ 167.4, 166.6, 166.1, 158.9, 139.0, 130.2, 116.9, 110.6, 60.4, 60.3, 54.5, 53.5, 38.9, 35.5, 28.3, 27.1, 22.2, 19.6, 14.3; HRMS-ESI (m/z) $[M+H]^+$ calcd. for $C_{19}H_{29}N_4O_4$: 377.2111 found: 377.2182

**Ethyl 4-hydroxy-2,6-dimethyl-1-((1-methyl-1*H*-pyrazol-4-yl)methyl)-2,3-dihydro-1 -pyrrolo[2,3-*b*]pyridine-5-carboxylate (8)**

To a solution of **7** (1.3 g, 3.45 mmol) in DMF (9 mL) was added sodium ethoxide (21% wt./EtOH, 3.88 mL, 10.37 mmol) and the mixture was placed in a pre-heated oil bath at 100 °C and stirred for 4 h. The reaction was cooled to 0 °C and the pH was adjusted by addition of 1 N HCl to pH

= 8-9. The solvents were removed under vacuum; the residue was dissolved in dichloromethane (3 x 20 mL) and washed with water. The organic layers were combined and dried over $MgSO_4$. The solvent was removed under vacuum. The residue was purified by flash chromatography using hexane/ethyl acetate (50/50 to 40/60) to afford the desired compound **8** (797 mg, 70%); $^1$H NMR (400 MHz, $CDCl_3$) *δ* 11.86 (s, 1H), 7.36 (s, 1H), 7.25 (s, 1H), 4.91 (d, *J* = 15.3 Hz, 1H), 4.35 (q, *J* = 7.0 Hz, 2H), 4.04 (d, *J* = 15.4 Hz, 1H), 3.82 (s, 3H), 3.80 – 3.74 (m, 1H), 3.08 – 3.01 (m, 1H), 2.65 (s, 3H), 2.43 (ddd, *J* = 15.7, 7.0, 0.9 Hz, 1H), 1.39 (t, *J* = 7.0 Hz, 3H), 1.27 (d, *J* = 7.0 Hz, 3H); $^{13}$C NMR (101 MHz, $CDCl_3$) *δ* 171.8, 164.4, 163.4, 162.3, 139.3, 129.5, 117.4, 101.7, 101.45, 61.2, 55.3, 39.0, 35.8, 31.1, 27.8, 19.8, 14.4; HRMS-ESI (m/z) $[M+H]^+$ calcd. for $C_{17}H_{23}N_4O_3$: 331.1692, found: 331.1764.

**Ethyl 4-hydroxy-2,6-dimethyl-1-((1-methyl-1*H*-pyrazol-4-yl)methyl)-1*H*-pyrrolo[2,3-*b*]pyridine-5-carboxylate (9)**

A solution of **8** (1 g, 3.02 mmol) in toluene (15 mL) and 2,3-dichloro-5,6-dicyano-1,4-benzoquinone (DDQ) (1.03 g, 4.5 mmol) was stirred at rt for 20 min. The solvent was removed under reduced pressure. The residue was purified by flash chromatography using hexane/ethyl acetate (50/50 to 40/60) to afford the desired compound **9** (575 mg , 58%); $^1$H NMR (400 MHz, $CDCl_3$-*d*) *δ* 12.61 (s, 1H), 7.39 (s, 1H), 7.16 (s, 1H), 6.31 (s, 1H), 5.26 (s, 2H), 4.46 (q, *J* = 7.1 Hz, 2H), 3.80 (s, 3H), 2.83 (s, 3H), 2.38 (s, 3H), 1.46 (t, *J* = 7.1 Hz, 3H); $^{13}$C NMR (101 MHz, $CDCl_3$) δ 172.8, 163.1, 155.2, 138.3, 134.5, 129.1, 118.8, 107.34, 101.8, 97.1, 61.6, 39.1, 36.0, 28.1, 14.4, 13.1; HRMS-ESI (m/z) $[M+H]^+$ calcd. for $C_{17}H_{21}N_4O_3$: 329.1535, found: 329.1605.

**Ethyl 2,6-dimethyl-1-((1-methyl-1*H*-pyrazol-4-yl)methyl)-4-(((trifluoromethyl)sulfonyl) oxy)-1H-pyrrolo[2,3-*b*]pyridine-5-carboxylate (10)**

To a solution of **9** (356 mg, 1.08 mmol) in dichloromethane (4 mL) was added triethylamine (0.218 ml, 1.62 mmol) at 0 °C followed by dropwise addition of $Tf_2O$ (0.194 mL, 1.18 mmol). The mixture was stirred at rt for 1h before being concentrated under vacuum. The crude product was purified by flash chromatography using hexane/ethyl acetate (50/50) to afford the desired compound **10** (497 g, 90 %); $^{1}$H NMR (400 MHz, $CDCl_3$) $\delta$ 7.39 (s, 1H), 7.19 (s, 1H), 6.28 (s, 1H), 5.29 (s, 2H), 4.43 (q, *J* = 7.1 Hz, 2H), 3.80 (s, 3H), 2.75 (s, 3H), 2.45 (s, 3H), 1.41 (t, *J* = 7.1 Hz, 3H); $^{13}$C NMR (101 MHz, $CDCl_3$) $\delta$ 165.8, 151.5, 150.3, 145.2, 139.6, 138.3, 129.2, 118.6 (q, *J* = 320.5 Hz), 117.9, 114.5, 111.0, 96.1, 62.2, 39.1, 36.4, 24.1, 14.1, 13.3; $^{19}$F NMR (377 MHz, $CDCl_3$) $\delta$ -73.45; HRMS-ESI (m/z) $[M+H]^+$ calcd. for $C_{18}H_{20}F_3N_4O_5S$: 461.1028, found: 461.1151.

**Ethyl 4-(4-chlorophenyl)-2,6-dimethyl-1-((1-methyl-1*H*-pyrazol-4-yl)methyl)-1*H*-pyrrolo[2,3-*b*]pyridine-5-carboxylate (11)**

To a solution of **10** (500 mg, 1.08 mmol) in dry DMF (5 mL) were added 4-chlorophenylboronic acid (194 mg, 1.24 mmol) and potassium carbonate (447 mg, 3.24 mmol), The reaction mixture was degassed with a stream of nitrogen for 5 min and then tetrakis-triphenylphosphine palladium ($Pd(PPh_3)_4$) (125 mg, 0.108 mmol) was added thereto. The reaction was degassed one more time with a stream of nitrogen, and heated under nitrogen atmosphere for 4 h at 100°C. The reaction solution was cooled to room temperature. The insoluble substances were removed by filtration through a celite pad. The filtrate was concentrated under reduced pressure, and then the residue was purified using silica gel column chromatography hexane-ethyl acetate (60/40 to 50/50) to give target compound **11** (364 mg, 80%); $^{1}$H NMR (400 MHz, $CDCl_3$) $\delta$ 7.47 – 7.33 (m, 5H), 7.20 (s, 1H), 6.02 (s, 1H), 5.32 (s, 2H), 4.10 (q, *J* = 7.1 Hz, 2H), 3.79 (s, 3H), 2.72 (s, 3H), 2.39 (s, 3H), 1.02 (t, *J* = 7.1 Hz, 3H); $^{13}$C NMR (101 MHz, $CDCl_3$) $\delta$ 170.1, 148.4, 147.7, 138.3, 138.2, 137.8, 136.5, 134.0, 130.0, 129.1, 128.6, 121.0, 118.6, 117.1, 97.9, 61.1, 39.0, 36.0, 23.3, 13.8, 13.1; HRMS-ESI (m/z) $[M+H]^+$ calcd. for $C_{23}H_{24}ClN_4O_2$: 423.1510, found: 423.1582.

**(4-(4-Chlorophenyl)-2,6-dimethyl-1-((1-methyl-1*H*-pyrazol-4-yl)methyl)-1*H*-pyrrolo[2,3-*b*]pyridin-5-yl)methanol (12)**

To a solution of compound **11** (370 mg, 0.87 mmol) in anhydrous dichloromethane (10 mL) under nitrogen atmosphere was added a 1 M DIBAL/hexane solution (5 mL, 5 mmol) over 5 min at -78°C. The resulting solution was stirred for 2 h at the same temperature before slow addition of a saturated aqueous solution of $NH_4Cl$ (10 mL). The mixture was then diluted with dichloromethane (4 mL) and stirred for 30 min while slowly raising the temperature to rt. The mixture was filtered over celite. The cake was washed twice with dichloromethane (10 mL). The organic layer was separated, and the aqueous layer was extracted with dichloromethane (10 mL x2). The organic layers were combined, washed with brine (10 mL) and dried with anhydrous $MgSO_4$, and the solvent was concentrated under reduced pressure. The residue was purified using silica gel column chromatography hexane/ethyl acetate (60/40 to 50/50) to give compound **12** (270 mg, 82%). $^1$H NMR (400 MHz, MeOD) *δ* 7.45 – 7.39 (m, 4H), 7.38 (d, *J* = 0.7 Hz, 1H), 7.20 (s, 1H), 5.87 (s, 1H), 5.30 (s, 2H), 4.62 (s, 2H), 3.76 (s, 3H), 2.79 (s, 3H), 2.36 (s, 3H); $^{13}$C NMR (101 MHz, MeOD) *δ* 151.5, 146.9, 140.6, 138.2, 136.5, 136.3, 133.7, 130.9, 129.2, 128.4, 122.9, 118.9, 118.2, 97.9, 59.6, 39.0, 35.8, 22.8, 22.7, 13.1, HRMS-ESI (m/z) $[M+H]^+$ calcd. for $C_{21}H_{22}ClN_4O$: 381.1404, found: 381.1472.

**4-(4-Chlorophenyl)-2,6-dimethyl-1-((1-methyl-1*H*-pyrazol-4-yl)methyl)-1*H*-pyrrolo[2,3-*b*]pyridine-5-carbaldehyde (13)**

To a solution of compound **12** (0.31 g, 0.82 mmol) in anhydrous dichloromethane (5 mL) was added pyridinium chlorochromate (PCC) (0.35 g, 1.63 mmol) and silica gel (0.245 g, 70% by weight of PCC) at 0 °C, under nitrogen atmosphere. The resulting mixture was stirred at rt for 3 h and the reaction was filtered. The residue was washed with dichloromethane (10 mL) and the filtrate was concentrated under reduced pressure. The residue was purified using silica gel column chromatography dichloromethane /methanol (99/1 to 95/5) to give compound **13** (0.175 g, 56%); $^1$H NMR (400 MHz, $CDCl_3$) δ 10.06 (s, 1H), 7.47 – 7.43 (m, 3H), 7.38 – 7.32 (m, 2H), 7.29 (s, 1H), 6.02 (s, 1H), 5.35 (s, 2H), 3.81 (s, 3H), 2.95 (s, 3H), 2.41 (m, 3H). $^{13}$C NMR (101 MHz, $CDCl_3$) *δ* 192.8, 153.9, 148.0, 145.7, 138.4, 138.3, 134.6, 134.1, 131.3, 129.2, 128.6,

121.0, 118.2, 118.1, 98.6, 39.0, 36.0, 25.5, 13.1; HRMS-ESI (m/z) $[M+H]^+$ calcd. for $C_{21}H_{20}ClN_4O$: 379.1247, found: 379.1319.

**2-(4-(4-Chlorophenyl)-2,6-dimethyl-1-((1-methyl-1*H*-pyrazol-4-yl)methyl)-1*H*-pyrrolo[2,3-*b*]pyridin-5-yl)-2-((trimethylsilyl)oxy)acetonitrile (14)**

To a solution of compound **13** (0.175 g, 0.46 mmol) in dichloromethane (5 mL) was added $ZnI_2$ (0.19 g, 0.60 mmol) at 0 °C under nitrogen atmosphere followed by addition of trimethylsilyl cyanide (0.23 mL, 1.85 mmol) over 5 min. The resulting mixture was stirred for 1 h at 0 °C and 3 h at 25 °C. The reaction was diluted with dichloromethane (10 mL), washed with water (10 mL), dried over anhydrous $MgSO_4$, and the solvent was concentrated under reduced pressure to give the target compound **14** (0.221 g, 100% crude yield) which was used in the next reaction without further purification.

**Methyl 2-(4-(4-chlorophenyl)-2,6-dimethyl-1-((1-methyl-1*H*-pyrazol-4-yl)methyl)-1*H*-pyrrolo[2,3-*b*]pyridin-5-yl)-2-hydroxyacetate (15)**

To a solution of compound **14** (0.2 g, 0.40 mmol) in methanol (10 mL) was added sulfuric acid (0.89 mL, 16.7 mmol) dropwise at 0 °C. The reaction mixture was then stirred at 75 °C for 24 h before being cooled to 0 °C and neutralized with 2N NaOH to pH 7.5. Solvents were removed under reduced pressure. The mixture was diluted with ethyl acetate (10 mL) and washed with water (10 mL). The aqueous phase was extracted with ethyl acetate (5 mL x 2) and the combined organic layers were dried over $Na_2SO_4$ and concentrated under vacuum. The resulting residue was purified by silica gel column chromatography using hexane/ethyl acetate (90/10 to 30/70) to provide **15** (0.050 g, 27%) along with starting material **14** (0.070 g); $^1H$ NMR (400 MHz, MeOD) δ 7.56 – 7.37 (m, 5H), 7.32 (s, 1H), 5.81 (s, $J$ = 1.1 Hz, 1H), 5.37 (s, 1H), 5.33 (s, 2H), 3.76 (s, 3H), 3.62 (s, 3H), 2.64 (s, 3H), 2.35 (s, 3H); $^{13}C$ NMR (101 MHz, MeOD) δ 175.7, 151.7, 147.8, 142.5, 139.0, 138.8, 137.4, 135.1, 132.6, 132.0, 130.8, 129.5, 123.8, 120.3, 119.8, 99.0,

70.2, 52.9, 38.8, 36.6, 22.9, 13.0; HRMS-ESI (m/z) $[M+H]^+$ calcd. for $C_{23}H_{24}ClN_4O_3$: 439.1459, found: 439.1470.

**Methyl 2-(4-(4-chlorophenyl)-2,6-dimethyl-1-((1-methyl-1*H*-pyrazol-4-yl)methyl)-1*H*-pyrrolo[2,3-*b*]pyridin-5-yl)-2-oxoacetate (16)**

To a solution of compound **15** (0.050 g, 0.114 mmol) in dichloromethane (DCM) (5 mL) at 0 °C was added in two portions Dess-Martin periodinane (DMP) (0.063 g, 0.148 mmol). The resulting mixture was stirred at rt for 2 h before being concentrated under vacuum and purified by chromatography using hexane/ethyl acetate (90/10 to 40/60) to provide **16** (0.043 g, 87% yield); $^1H$ NMR (400 MHz, $CDCl_3$) δ 7.47 – 7.41 (m, 3H), 7.37 – 7.31 (m, 2H), 7.27 (s, 1H), 6.08 (d, *J* = 1.1 Hz, 1H), 5.35 (s, 2H), 3.83 (s, 3H), 3.39 (s, 3H), 2.74 (s, 3H), 2.42 (d, *J* = 1.1 Hz, 3H); $^{13}C$ NMR (101 MHz, $CDCl_3$) δ 190.2, 163.4, 151.5, 148.1, 140.7, 138.7, 138.4, 135.1, 135.0, 131.5, 129.4, 128.9, 122.4, 118.3, 117.3, 98.1, 52.7, 39.1, 36.1, 24.0, 13.3; HRMS-ESI (m/z) $[M+H]^+$ calcd. for $C_{23}H_{22}ClN_4O_3$: 437.1302, found: 437.1415.

**Methyl (*S*)-2-(4-(4-chlorophenyl)-2,6-dimethyl-1-((1-methyl-1*H*-pyrazol-4-yl)methyl)-1*H*-pyrrolo[2,3-*b*]pyridin-5-yl)-2-hydroxyacetate (17)**

To a solution of compound **16** (0.043 g, 0.098 mmol) in toluene (4 mL) was added (*R*)-1-methyl-3,3-diphenylhexahydropyrrolo[1,2-*c*][1,3,2]oxazaborole ((*R*)-Me-CBS, 0.024 mL, 0.024 mmol, 1 M in toluene) under nitrogen atmosphere. The mixture was cooled to -35 °C (EtOH/dry ice) and then catecholborane (0.30 mL, 0.295 mmol, 1 M in THF) was slowly added over ~30 min. The mixture was kept at -35 °C for 30 min and then allowed to warm to 0 °C in ~ 2 h. 2M $Na_2CO_3$/water (4 mL) was added followed by EtOAc (10 mL) and the mixture was stirred for 5 min at rt. Water was added and the organic phase was washed with 1 M NaOH/water (4 mL), dried over $Na_2SO_4$, and concentrated. The residue was purified using silica gel column chromatography hexane/ethyl acetate (90/10 to 30/70) to provide compound **17** (0.020 g, 46%). $^1H$ NMR (400 MHz, MeOD) *δ* 7.58 – 7.38 (m, 5H), 7.33 (s, 1H), 5.83 (d, *J* = 1.1 Hz, 1H), 5.36 (s,

3H), 3.79 (s, 3H), 3.65 (s, 3H), 2.64 (s, 3H), 2.39 (d, *J* = 1.0 Hz, 3H); $^{13}$C NMR (101 MHz, MeOD) *δ* 175.7, 151.7, 147.9, 142.5, 139.0, 138.8, 137.4, 135.2, 132.6, 132.0, 130.8, 129.5, 123.8, 120.4, 119.9, 99.0, 70.2, 52.9, 38.8, 36.6, 22.9, 13.0; HRMS-ESI (m/z) $[M+H]^+$ calcd. for $C_{23}H_{24}ClN_4O_3$: 439.1459, found 439.1535.

**Methyl (*S*)-2-(*tert*-butoxy)-2-(4-(4-chlorophenyl)-2,6-dimethyl-1-((1-methyl-1*H*-pyrazol-4-yl)methyl)-1*H*-pyrrolo[2,3-*b*]pyridin-5-yl)acetate (18)**

To a solution of **17** (0.020 g, 0.045 mmol) in tert-butyl acetate (0.489 mL, 3.65 mmol) was added dropwise perchloric acid (15 μL, 0.182 mmol) and the mixture was stirred at rt for 3 h. Saturated $NaHCO_3$/water (3 mL) was added slowly and then extracted with ethyl acetate (10 mL). The organic phase was washed with brine, dried, concentrated and purified by chromatography using hexane/ethyl acetate (90/10 to 40/60) to provide compound **18** (0.016 g, 72%); $^1$H NMR (400 MHz, MeOD) *δ* 7.58 – 7.51 (m, 3H), 7.46 – 7.41 (m, 2H), 7.35 (s, 1H), 5.84 (d, *J* = 1.2 Hz, 1H), 5.36 (d, *J* = 2.2 Hz, 2H), 5.34 (s, 1H), 3.79 (s, 3H), 3.76 (s, 3H), 2.65 (s, 3H), 2.39 (d, *J* = 1.0 Hz, 3H), 0.92 (s, 9H); $^{13}$C NMR (101 MHz, MeOD) *δ* 175.7, 152.1, 147.7, 141.1, 139.1, 138.7, 137.6, 135.3, 132.8, 132.7, 130.9, 129.6, 129.5, 124.6, 120.4, 119.6, 98.9, 76.8, 71.2, 52.9, 38.8, 36.6, 28.3, 23.6, 13.0 ; HRMS-ESI (m/z) $[M+H]^+$ calcd. for $C_{27}H_{31}ClN_4O_3$: 495.2085, found: 495.2159.

**(*S*)-2-(*tert*-Butoxy)-2-(4-(4-chlorophenyl)-2,6-dimethyl-1-((1-methyl-1*H*-pyrazol-4-yl)methyl)-1*H*-pyrrolo[2,3-*b*]pyridin-5-yl)acetic acid (19)**

To a solution of compound **18** (0.016, 0.032 mmol) in THF (3 mL) was added 4N NaOH (24 μL) and methanol (24 μL). The mixture was stirred for 18 h at 25 °C and 4N HCl (24 μL) was added to neutralize the mixture. The resulting mixture was concentrated under vacuum and the residue was purified by column chromatography using dichloromethane/methanol (99/ 1 to 85/15) to give compound **19** (0.010 g, 66%); $^1$H NMR (400 MHz, MeOD) *δ* 7.70 (m, 1H), 7.57 – 7.50 (m, 2H), 7.48 – 7.43 (m, 1H), 7.42 (s, 1H), 7.34 (d, *J* = 0.8 Hz, 1H), 5.86 (d, *J* = 1.1 Hz, 1H), 5.36 (d, *J* = 4.8 Hz, 2H), 5.31 (s, 1H), 3.78 (s, 3H), 2.71 (s, 3H), 2.38 (d, *J* = 1.0 Hz, 3H), 0.91 (s, 9H); $^{13}$C

NMR (101 MHz, MeOD) $\delta$ 152.2, 147.7, 141.1, 139.1, 138.5, 137.7, 135.3, 133.0, 132.7, 130.9, 129.6, 129.4, 125.2, 120.4, 119.5, 99.0, 76.8, 71.0, 38.8, 36.6, 28.3, 23.7, 13.0 ; HRMS-ESI (m/z) $[M+H]^+$ calcd. for $C_{27}H_{31}ClN_4O_3$: 481.1928, found: 481.1999.
